# Convergent antibody responses to the SARS-CoV-2 spike protein in convalescent and vaccinated individuals

**DOI:** 10.1101/2021.05.02.442326

**Authors:** Elaine C. Chen, Pavlo Gilchuk, Seth J. Zost, Naveenchandra Suryadevara, Emma S. Winkler, Carly R. Cabel, Elad Binshtein, Rachel E. Sutton, Jessica Rodriguez, Samuel Day, Luke Myers, Andrew Trivette, Jazmean K. Williams, Edgar Davidson, Shuaizhi Li, Benjamin J. Doranz, Samuel K. Campos, Robert H. Carnahan, Curtis A. Thorne, Michael S. Diamond, James E. Crowe

## Abstract

Unrelated individuals can produce genetically similar clones of antibodies, known as public clonotypes, which have been seen in responses to different infectious diseases as well as healthy individuals. Here we identify 37 public clonotypes in memory B cells from convalescent survivors of SARS-CoV-2 infection or in plasmablasts from an individual after vaccination with mRNA-encoded spike protein. We identified 29 public clonotypes, including clones recognizing the receptor-binding domain (RBD) in the spike protein S1 subunit (including a neutralizing, ACE2-blocking clone that protects *in vivo*), and others recognizing non-RBD epitopes that bound the heptad repeat 1 region of the S2 domain. Germline-revertant forms of some public clonotypes bound efficiently to spike protein, suggesting these common germline-encoded antibodies are preconfigured for avid recognition. Identification of large numbers of public clonotypes provides insight into the molecular basis of efficacy of SARS-CoV-2 vaccines and sheds light on the immune pressures driving the selection of common viral escape mutants.

## Introduction

Severe acute respiratory syndrome coronavirus 2 (SARS-CoV-2) is the causative agent of COVID-19 and the ongoing worldwide pandemic. SARS-CoV-2 is a betacoronavirus, with other virus family members having caused global outbreaks including the 2003 SARS-CoV-1 and 2012 Middle East Respiratory Syndrome coronavirus (MERS-CoV) epidemics. The spike (S) protein is the principal antigen recognized by the protective antibody response against SARS-CoV-2^1,2^. The S protein is cleaved into S1, which includes the receptor-binding domain (RBD) and the N-terminal domain (NTD), and S2, which contains the fusion peptide and heptad repeats HR1 and HR2 and mediates fusion between virus and host cell membrane^3,4^. SARS-CoV-2 and SARS-CoV-1 share approximately 80% amino acid sequence identity, and both use human angiotensin-converting enzyme 2 (ACE2) as an entry receptor through binding mediated by the RBD^5–7^.

Monoclonal antibodies (mAbs) targeting the SARS-CoV-2 S protein have been a focus for development of medical countermeasures against COVID-19. Many studies have identified antibodies to the S1 and S2 regions on the S protein, with the majority of neutralizing antibodies targeting the RBD in S1 and inhibiting ACE2 binding^8–12^. Multiple RBD-specific mAbs have been developed as monotherapies or cocktail therapeutics, and two (Lilly mAbs bamlanivimab [LY-CoV555] and etesevimab [LY-CoV016, also known as JS016] as well as Regeneron mAbs casirivimab and imdevimab) have received Emergency Use Authorization (EUA)^13,14^. Additionally, multiple vaccines eliciting antibodies to the S protein are being deployed globally under similar EUA^15–17^.

In recent years, public B cell clonotypes have been identified in the human antibody repertoires formed in response to diverse viruses including Ebola^18–20^, influenza^21–25^, human immunodeficiency virus 1 (HIV-1)^26–29^, hepatitis C^30,31^, SARS-CoV-2^32–34^, and in healthy individuals^35,36^. These studies reveal a convergence of B cell selection resulting in circulating B cells clones with genetically similar antigen receptor genes in multiple individuals. The selection of public B cell clonotypes often has a structural basis mediated by low-affinity recognition of virus surface antigens by unmutated germline-encoded naïve B cell receptors that are preconfigured for binding and cell activation. Public clonotypes are of great interest, since the understanding of viral epitopes that commonly induce antibodies in humans has implications for predicting the most common responses to vaccines in large populations. With newer single-cell technologies, it is now possible to obtain paired heavy and light chain antibody variable gene sequences, allowing investigators to describe gene usage and study the function of recombinant antibodies expressed from synthesized cDNA in a large scale. This approach is powerful, since coupling genotype with function allows analysis of the role of public B cell clonotypes in the response to infection or vaccination.

There have been several efforts to characterize public clonotypes in the response to SARS-CoV-2, with most work focused on neutralizing public clonotypes^9,32,33,37^ that target the S1 domain of the S trimer, more specifically the RBD and NTD domains. However, it is less clear if public clonotypes are directed to other sites on the S trimer such as the S2 domain. Epitopes on the S2 domain may be of interest, as these sites may be more conserved than those in RBD in different strains of coronavirus due to functional constraints associated with the viral fusion mechanism. This sequence conservation reflects the fact that the S2 domain contains the HR1, HR2, and fusion loop of the S trimer, all of which are required for viral entry in coronaviruses. Given the importance of defining immune responses to SARS-CoV-2 infection or vaccination, we sought to identify the spectrum of public clonotypes, including less well studied ones directed to non-RBD regions or those lacking neutralizing activity. Understanding public clonotype recognition to all antigenic domains of the S trimer, and not just the RBD, delineates the B cell response to SARS-CoV-2 to regions that are more conserved in the S protein. In this study, we identified 37 total public clonotypes, 27 of which are shared between vaccinated and convalescent individuals. Of the public clones identified a detailed analysis of three public clonotypes (Groups 1, 2, and 3) not previously described and comparisons of public antibodies discovered from large-scale discovery efforts were investigated. We found that shared clonotypes comprise a substantial proportion of the elicited human B cell response to the S trimer. We also compared the response following infection or mRNA vaccination to investigate the genetic basis for the efficacy of mRNA vaccines in the population. These data show that many clonotypes are shared between convalescent and vaccinated individuals. Finally, as if diverse individuals independently make the same antibody in response to an antigen, it induces selective pressure on that epitope. And therefore, the frequent occurrence of public clonotypes recognizing sites of vulnerability on S protein that tolerate mutations may explain the rapid emergence of particular SARS-CoV-2 variant viruses in the field. The collective immunity mediated by the large number of public clonotypes described here on particular sites of vulnerability like drive the independent escape events leading to emergence of variants of concern in diverse geographic areas.

## RESULTS

### Identification of public clonotypes

To identify a comprehensive set of public clonotypes in the B cell response to SARS-CoV-2, we first collected antibody variable gene sequences for SARS-CoV-2 human mAbs from existing publications that had isolated mAbs from individuals with a history of SARS-CoV-2 infection^8–12,38–41^. This search identified a panel of 2,865 paired heavy and light chain variable gene sequences for analysis. We clustered all sequences by binning the clones based on the inferred immunoglobulin heavy variable (*IGHV*) gene, immunoglobulin heavy joining (*IGHJ*) gene, and the amino acid length of the heavy chain complementarity determining region 3 (CDRH3). These sequences then were clustered according to 70% nucleotide sequence identity in the DNA sequence encoding the CDRH3. Next, the sequences were binned further based on the inferred immunoglobulin light variable gene (*IGLV* or *IGKV*) and immunoglobulin light joining (*IGLJ* or *IGKJ*) genes. Clusters meeting these similarity criteria in both heavy and light chains with sequences originating from two or more individuals were deemed public clonotypes (**Extended Data Fig.1**). Eleven public clonotypes were identified in the repertoires of subjects with prior natural infection (**Fig. 1a, b, c**), and these clones are encoded by a variety of heavy and light chain variable genes. Of the 11 public clonotypes identified, five of the heavy chain genes have been reported previously to encode potently neutralizing SARS-CoV-2 antibodies that bind to the RBD: *IGHV3-53*^*32*^*, IGHV1-58*^*33*^*, IGHV3-30, IGHV3-30-3*^*9*^*, IGHV3-66*^*32*^, whereas three have not been reported: *IGHV1-69, IGHV4-59, IGHV3-7*. *IGHV3-53* and *IGHV3-66* are commonly observed in antibodies in SARS-CoV-2 patients^9^ since the germline gene segments encode amino acid motifs that are preconfigured for RBD binding^32^. *IGHV1-58* also commonly encodes antibodies that neutralize SARS-CoV-2, as this germline gene segment encodes motifs that mediate binding to the S protein^33^. Notably, *IGHV1-58* encodes the mAb COV2-2196, which is the basis for one of the two antibodies in a cocktail currently in Phase III clinical trials^33,42^. Clonally expanded B cell populations containing potently neutralizing antibodies encoded by *IGHV3-30* also have been found in multiple individuals^9^. However, the role of *IGHV1-69*, *IGHV4-59*, and *IGHV3-7* public clonotypes in SARS-CoV-2 responses remains unknown. In this paper, for clarity, we designated public clonotypes incorporating these additional three V_H_ gene segments as members of Group 1, 2, or 3 mAbs, respectively (**Fig. 1c, d, e**). Group 1 is shared by two donors from the cohort we studied and includes mAbs COV2-2002 and COV2-2333. Group 2 is shared by a donor from our group and a previously described donor IDCnC2^39^ and includes antibodies COV2-2164 and CnC2t1p1_B10. Lastly, Group 3 is shared by a donor from our group and a previously described donor COV107^9^ and includes antibodies COV2-2531 and C126 (**Fig. 1c, d, e**). cDNAs for the antibody variable genes encoding each of the six antibodies from the three groups of public clonotypes were synthesized and cloned into an IgG1 expression vector, as previously described^43^.

**Figure 1.**
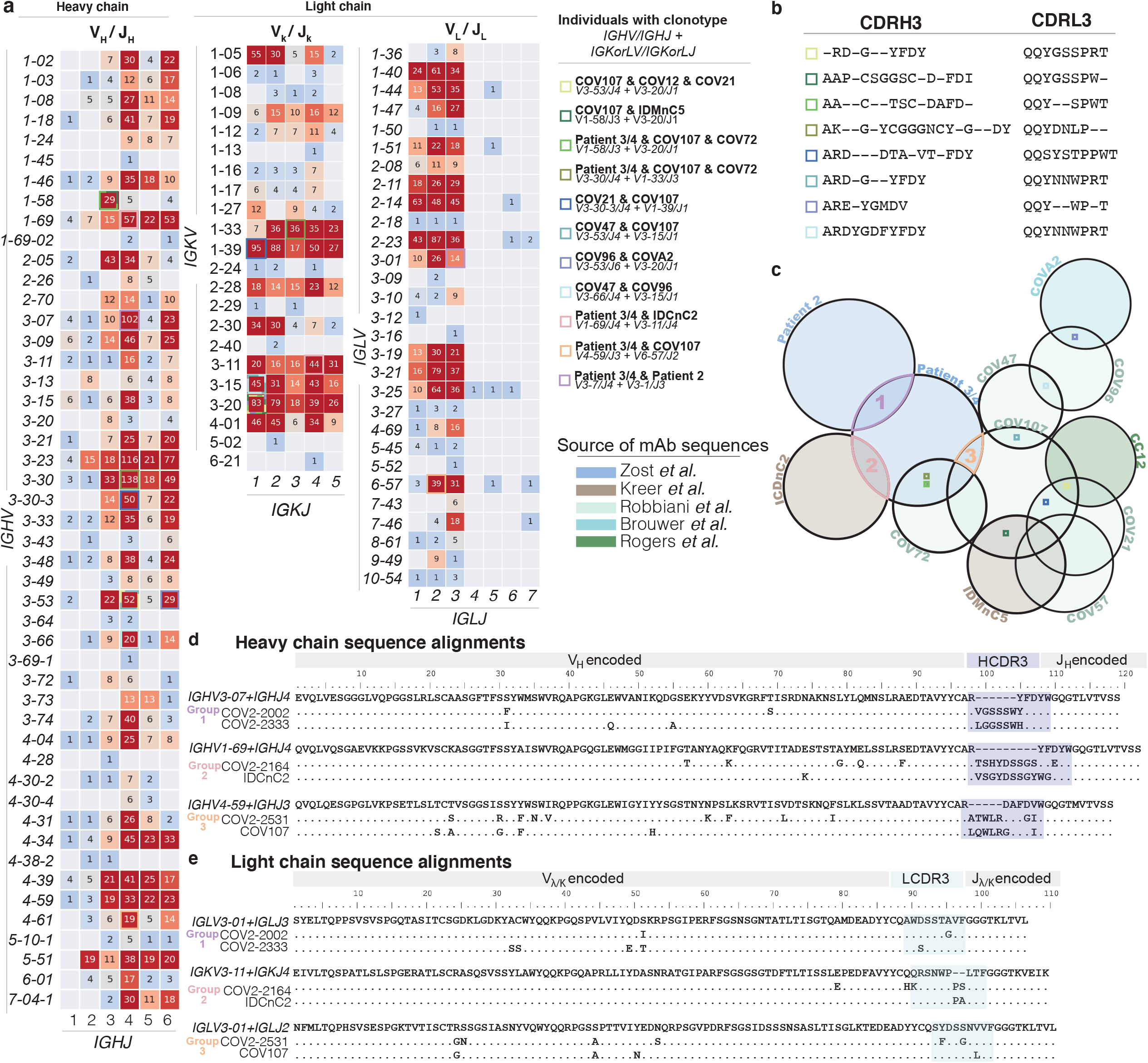
Sequence characteristics of monoclonal antibodies to SARS-CoV-2. **a.** Available sequences of mAbs to SARS-CoV-2 from multiple publications were obtained from public databases. Numbers inside each box represents the number of sequences with the indicated gene usage. Colored boxes represent public clonotypes that are shared between the individuals listed. **b.** CDR3 sequences of the heavy and light chains of each of the remaining eight public clonotypes are shown. Dashes represent amino acids that differed in the public clonotype. Each box color correlates to the public clonotypes in **Fig 1c**. **c.** A Venn diagram illustrating all of the public clonotypes identified between naturally-infected individuals. The colored boxes in the Venn diagram overlaps represent the public clonotypes identified in **Fig 1a**. Novel public clonotypes, designated as Groups 1, 2, or 3, are highlighted in the purple, pink, or orange overlaps respectively. **d.** Multiple sequence alignments of the heavy chain sequences for Groups 1, 2, or 3 to their respective inferred germline genes *IGHV 3-07/IGHJ4, IGHV1-69/IGHJ4, or IGHV4-59/IGHJ3.* The CDRH3 sequence is highlighted in dark blue. **e.** Multiple sequence alignments of the light chain sequences for Groups 1, 2, or 3 to their respective inferred germline genes *IGLV3-01/IGLJ3, IGKV3-11/IGKJ4, or IGHV3-01/IGLJ2.* The CDRL3 sequence is highlighted in light blue.

### Functional properties of identified public clonotype antibodies

To examine the binding properties of antibodies in these three new SARS-CoV-2 public clonotypes, we tested six recombinant purified antibodies, two for each public clonotype, for binding to recombinant stabilized trimeric prefusion ectodomain of the SARS-CoV-2 S protein (S6P_ecto_), SARS-CoV-2 RBD, or recombinant stabilized trimeric prefusion ectodomain of the SARS-CoV-1 S protein (S2P_ecto_) proteins by ELISA (**Extended Data Fig. 2**). The two Group 1 antibodies, COV2-2002 and COV2-2333, did not bind to SARS-CoV-2 RBD, but both bound to SARS-CoV-2 S6P_ecto_ and SARS-CoV-1 S2P_ecto_ proteins. However, they did not saturate in binding to SARS-CoV-1 S2P_ecto_ at the maximum concentration tested (400 ng/mL) indicating relatively weak binding to recombinant SARS-CoV-1 S2P_ecto_. Group 2 antibodies, which include COV2-2164 and CnC2t1p1_B10, did not bind to SARS-CoV-2 RBD, but both bound to SARS-CoV-2 S6P_ecto_ and SARS-CoV-1 S2P_ecto_ proteins. Group 3 antibodies, which include COV2-2531 and C126, bound to SARS-CoV-2 S6P_ecto_ and SARS-CoV-2 RBD proteins (**Fig. 2a, h**). However, antibodies from Group 3 did not bind SARS-CoV-1 S2P_ecto_.

**Figure 2.**
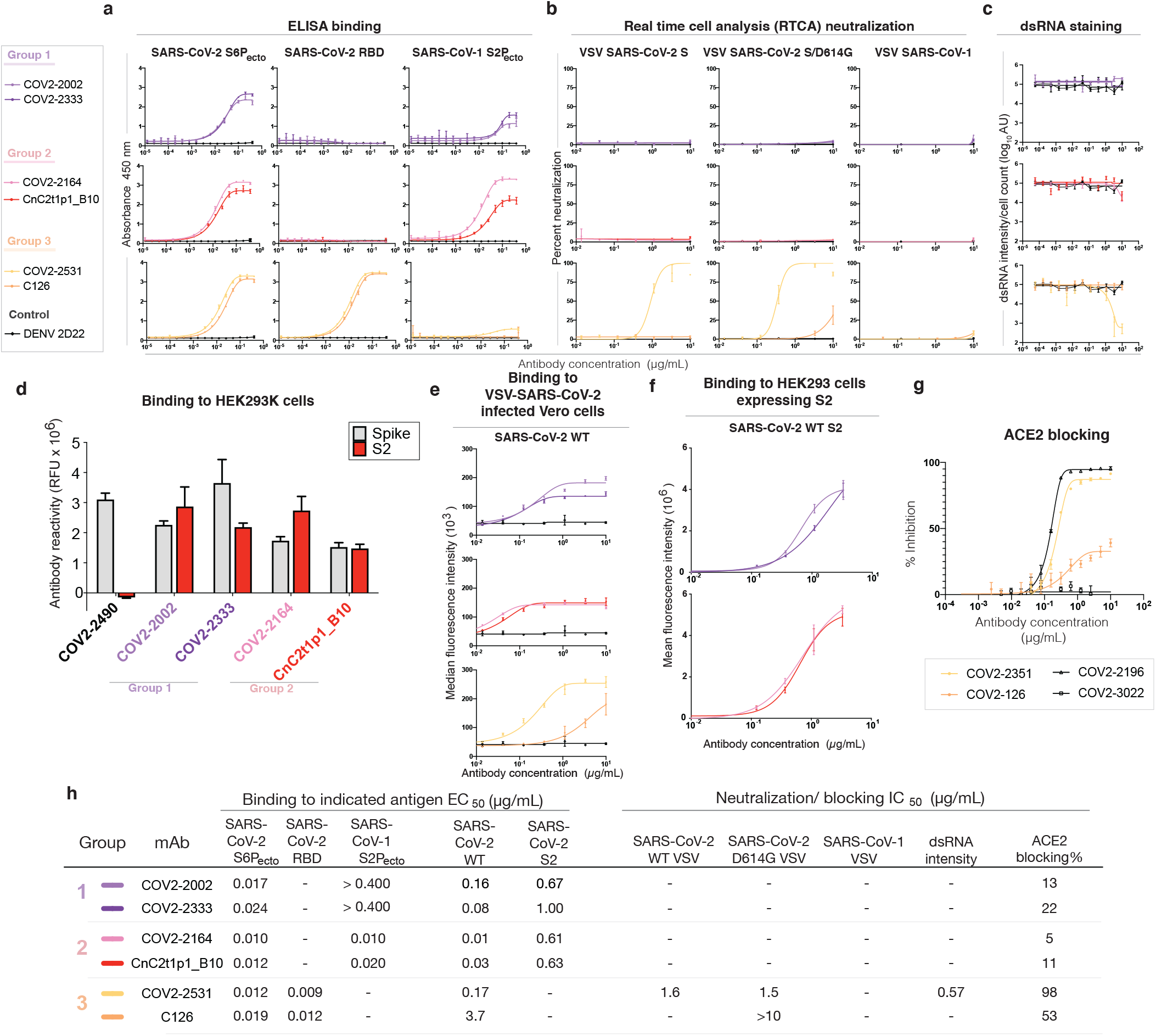
Reactivity and functional activity of Groups 1, 2, and 3 antibodies. Group 1 antibodies are shown in light or dark purple, Group 2 antibodies are in red or pink, and Group 3 antibodies are in light or dark orange. MAb DENV 2D22 was used as a negative control antibody, as shown in the lines in black. All experiments are performed in biological replicates and technical triplicates. Biological replicate from representative single experiment shown. a. ELISA binding to SARS-CoV-2 S6P_ecto_, SARS-CoV-2 RBD, or SARS-CoV-1 S2P_ecto_ was measured by absorbance at 450 nm. Antibody concentrations starting at 0.4 µg/mL were used and titrated two-fold. b. Neutralization activity of antibodies to VSV-SARS-CoV-2, VSV-SARS-CoV-2/D614G, and VSV-SARS-CoV-1 determined by using real time cell analysis (RTCA) assay. The percent of neutralization is reported. Antibody concentrations started at 10 µg/mL and were titrated three-fold. c. Neutralization activity of antibodies to authentic SARS-CoV-2 (USA-WA1/2020) determined by measuring dsRNA intensity per cell count after Calu3 lung epithelial cells were inoculated with SARS-CoV-2. Antibody concentrations started at 10 µg/mL and were titrated three-fold. d. Antibody binding to full-length S (grey) or S protein C-terminus S2 region (red) expressed on the surface of HEK-293T cells that were fixed and permeabilized. Antibodies were screened at 1 µg/mL. Antibody reactivity was measured by flow cytometry and cellular florescence values were determined. COV2-2490, an NTD-directed antibody, was used as a control. e. Binding to VSV-SARS-CoV-2-infected Vero cells (SARS-CoV-2 WT) was measured using flow cytometry and median florescence intensity values were determined for dose-response binding curves. Antibody was diluted 3-fold staring from 10 µg/mL f. Binding to S protein C-terminus S2 region expressed on HEK-293T cells (SARS-CoV-2 WT S2) was measured using flow cytometry and mean fluorescence intensity values were determined for dose-response binding curves. Antibody was diluted 3-fold starting from 10 µg/mL. g. Inhibition of ACE2 binding curves for COV2-2531 or C126. Antibody concentrations started at 10 µg/mL and were titrated 3-fold to identify ACE2 blocking curves. COV2-2531 is shown in light orange, and C126 is shown in dark orange. h. Binding EC_50_ and neutralization IC_50_ values for each of the assay curves in **Fig 3a, b, c, d, e**. All values are denoted as µg/mL. ACE2 blocking was determined by measuring amount of ACE2 with FLAG tag binding in the presence of each antibody, measured by binding of an anti-FLAG antibody. Percent blocking is shown, calculated by using ACE2 binding without antibody as 0% blocking.

As antibodies from Groups 1 and 2 did not bind the RBD but cross-reacted to both SARS-CoV-2 S6P and SARS-CoV-1 S2P, we hypothesized that they might bind the S2 domain of the S trimer; SARS-CoV-2 infection can elicit antibodies that recognize cross-reactive epitopes on the S2 domain^44^. Antibodies were tested for binding against the S2 domain of SARS-CoV-2 S expressed on HEK-293T cells. An NTD-directed antibody COV2-2490 was used as a control. This experiment showed that Group 1 and 2 antibodies bound to S2 in a dose-dependent manner (**Fig. 2d, f, h**), and revealed that public clonotypes can be elicited to the S2 domain of the S trimer.

Antibodies from each group then were tested for neutralizing activity using a previously described real-time cell analysis (RTCA) assay that measures cellular impedance^8,43^. We used recombinant, infectious vesicular stomatitis virus (VSV) expressing the S proteins from SARS-CoV-2 (WA1/2019 strain), SARS-CoV-2/D614G, or SARS-CoV-1 (Urbani strain) (**Extended Data Fig. 2**). In addition, we used authentic infectious SARS-CoV-2 (WA1/2019) virus and Calu3 (human lung epithelial adenocarcinoma) cell monolayer cultures, and neutralization was measured by staining for double-stranded RNA, which is produced in the cytoplasm in virus-infected cells (**Extended Data Fig. S3**). Group 3 mAb COV2-2531 neutralized SARS-CoV-2 (VSV-SARS-CoV-2, and VSV-SARS-CoV-2/D614G (**Fig. 2b, c, h**) and authentic SARS-CoV-2, but not SARS-CoV-1. In contrast, another Group 3 mAb, C126 partially neutralized SARS-CoV-2/D614G variant but did not neutralize the WT VSV-SARS-CoV-2, VSV-SARS-CoV-1, or authentic virus. Groups 1 and 2 antibodies did not exhibit neutralizing capacity for any of the viral strains tested.

As both Group 3 antibodies exhibited neutralizing capacity, we considered that they might bind to the RBD and block virus attachment to ACE2, a principal mechanism of inhibition by RBD-targeted antibodies against SARS-CoV-2^42,45^. We tested whether each antibody could block binding of soluble trimeric S protein to recombinant human ACE2 protein in an ELISA. Only Group 3 antibodies blocked binding to ACE2 (**Fig. 2h**). Similar to the pattern we observed for neutralization, COV2-2531 fully blocked ACE2 binding, whereas C126 partially blocked binding, with less than 50% inhibition at maximal effect (**Fig. 2g**). Therefore, it is likely that COV2-2531 neutralizes virus infection at least in part by blocking binding to ACE2.

### Binding sites of identified clonotype antibodies

We used negative stain electron microscopy (EM) to image Fab-SARS-CoV-2 S6P_ecto_ complexes. Even though all of the antibodies bound to S protein in ELISA as IgG1, only the Group 3 antibody Fabs COV2-2531 and C126 formed complexes visualized on EM grids, suggesting that some antibodies may require an IgG format for strong binding (**Fig. 3a**, **Extended Data Fig. 4**). Low-resolution 3D reconstructions for COV2-2531 and C126 showed that these two antibodies bind the side of the RBD and recognize the cryptic face of the RBD that is accessible only in the “open” position of the RBD in the context of the S trimer (**Fig. 3b, c**).

**Figure 3.**
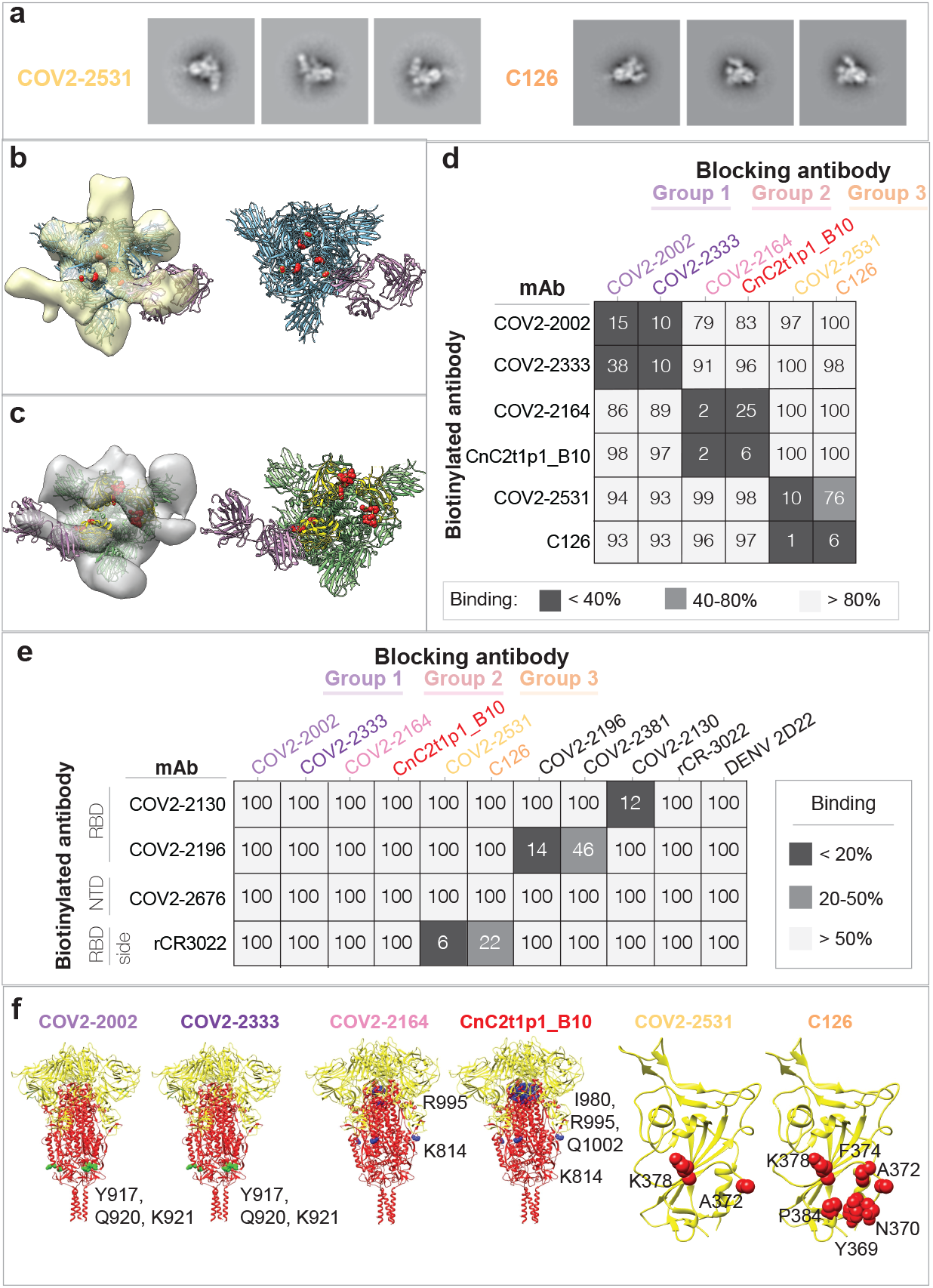
Epitope identification and structural characterization of antibodies. a. Negative stain EM of SARS-CoV-2 S6P_ecto_ protein in complex with Fab forms of different mAbs. Negative stain 2D classes of SARS-CoV-2 S protein incubated with COV2-2531 or C126. Box size is 128 pix at 4.36 Å/pix. b. MAb COV2-2531 3D volume with critical residues 372 and 275 shown in red on the S protein (blue). RBD is in the open position. Density corresponds to three fabs, as we docked a single Fab structure onto the EM density map, shown in magenta. c. MAb C126 3D volume with critical binding residues shown in red. The Fab is docked to a protomer of SARS-CoV-2 S protein in the open conformation. Top left is the RBD positioned in open conformation, with the other two protomers in the trimer in closed position. The S protein is shown in green, with the RBD in yellow. The Fab is shown in magenta. d. Competition-binding ELISA results for mAbs within each clonotype group. Unlabeled blocking antibodies applied to antigen first are listed across the top, while biotinylated antibodies that are added to antigen-coated wells second are indicated on the left. The number in each box represents percent un-competed binding of the biotinylated antibody in the presence of the indicated competing antibody. Heat map colors range from dark grey (<40% binding of the biotinylated antibody) to light grey (>80% binding of the biotinylated antibody). Experiment was performed in biological replicate and technical triplicate. Biological replicate from representative single experiment shown. e. Competition-binding ELISA data using Group 1, 2, or 3 antibodies against epitope-mapped reference antibodies. Biotinylated antibodies are indicated on the left, and the unlabeled antibodies applied to antigen first are indicated across the top. Heat map colors range from dark grey (<20% binding of the biotinylated antibody) to light grey (>50% binding of the biotinylated antibody). Experiment was performed in biological replicate and technical triplicates. Biological replicate from representative single experiment shown. f. Alanine scanning mutagenesis results for Group 1, 2 or 3 antibodies. S2 Epitope residues are shown (green spheres or blue spheres) on the S protein structure (PDB 6XR8), S1 is colored yellow, S2 red. RBD epitopes are shown in red on the RBD structure (PDB 6XR8). Primary data shown in **Fig. S5**.

We then tested binding of antibodies to the full-length membrane-bound S protein using infected Vero cells that were inoculated with VSV-SARS-CoV-2 chimeric viruses. We used a dengue virus specific antibody (DENV 2D22)^46^ and SARS-CoV-2-reactive antibody (COV2-2381)^8^ as controls (**Extended Data Fig. 5**). Antibodies from each of the groups bound to infected cells dose-dependently, with the Group 3 RBD-reactive antibodies exhibiting greater binding than the Group 1 or 2 S2-reactive antibodies (**Fig. 2e, h**). Binding of the Group 3 antibodies correlated with their neutralization capacity, as COV2-2531 showed greater binding than C126. The capacity to bind to infected cells also suggested that these antibodies could act *in vivo* not only by direct virus neutralization but also through Fc-mediated functions.

To identify if the antibodies within each discrete public clonotype group bind similar epitopes, we used competition-binding ELISA for pairwise comparison of antibodies binding to the S6P_ecto_ protein (**Fig. 3d**). As expected, members of each public clonotype group clustered with the other member of the same group by competition-binding. To begin to determine specific epitopes recognized by mAbs in each group, we competed the antibodies for binding against a larger group of epitope-mapped antibodies we previously described^8^, that covers various sites on the S protein, and against rCR3022^47^, which bind less well to the RBD of SARS-CoV-2 compared to SARS-CoV-1 and does not block ACE2 binding (**Fig. 2g**). Both Group 3 antibodies competed with rCR3022, with COV2-2531 exhibiting a higher level of competition than C126. None of the Group 1 or 2 antibodies competed with the reference antibodies tested (**Fig. 3e)**.

We then determined the critical binding residues at the amino acid level for each of the public clonotype antibodies by screening for binding to alanine-scanning mutant libraries of the SARS-CoV-2 S protein. Screening the RBD library revealed A372 and K378 as critical residues for COV2-2531 binding. For C126, we also identifiedA372 and K378, but with additional critical residues Y369, N370, F374, and P384 (**Fig. 3f**, **Extended data Fig. S6**). Notably, the identified residues are consistent with the binding site identified in the negative stain EM analyses and overlap with the epitope of CR3022^47^. It was curious that several SARS-CoV-2-specific neutralizing antibodies competed with CR3022, which also binds to SARS-CoV-1 but is non-neutralizing. It is of note that SARS-CoV-1 has an N-glycosylation site at N370, in the binding site for these two mAbs, which SARS-CoV-2 lacks^47^. This difference in glycosylation likely explains why COV2-2531 and C126 do not bind or neutralize SARS-CoV-1, even though they recognize the relatively conserved cryptic face of the RBD (**Extended Data Fig. S7**). In the alanine scanning libraries, native alanine residues are changed to serine. It is possible that A372 was identified as critical for binding by COV2-2531 and C126 because the A372S mutation results in the introduction of N-linked glycosylation of N370, rather than making direct side-chain contact with the antibodies.

Screening the Group 1 and 2 antibodies against the SARS-CoV-2 alanine scanning mutation library confirmed that they bound to the S2 domain. For the Group 1 antibodies COV2-2002 and COV2-2333, we identified critical residues for both antibodies (Y917, Q920, K921) in the heptad repeat (HR1) region of S2. These residues are conserved between SARS-CoV-2 and SARS-CoV-1. For the group 2 antibodies, screens identified two regions of residues that were specifically critical for binding. For both COV2-2164 and CnC2t1p1_B10, we identified K814 (also conserved between SARS-CoV-2 and SARS-CoV-1) as critical for binding. In addition, for both antibodies we also identified R995, and additionally for CnC2t1p1_B10, I980 and Q1002 (**Fig. 3f, Extended data Fig. S6**). K814 is not close to I980, R995, or Q1002 on the S protein structure. However, inspection of the available S protein structures (PDB: 6XR8 and 7C2L) suggested that residues I980, R995, and Q1002 are not readily accessible to antibodies in the full S protein, or even in the absence of S1. These residues make interactions that likely help maintain S2 structure, and so their mutation could indirectly affect Group 2 antibody binding. We conclude that K814 is an epitope residue for Group 1 antibodies, COV2-2002 and COV2-2333, as well as Group 2 antibodies, COV2-2164 and CnC2t1p1_B10. These results suggest that the mAbs in each public clonotype group have the essentially identical critical epitope residues.

### Functional properties of germline-revertant forms of antibodies from each identified public clonotype

To determine if the function of each antibody group was due to germline-encoded reactivity or the result of somatic mutations, we investigated the equivalent germline encoded antibodies. Heavy and light chain variable region sequences of antibodies COV2-2002, COV2-2164, and COV2-2531 were aligned with the germline sequences of [*IGHV3-7/IGHJ4*/ + *IGLV3-1/IGLJ3*], [*IGHV1-69*/IGHJ4 + *IGKV3-11/IGKJ4*], or [*IGHV4-59/IGHJ3* + *IGLV6-57/IGLJ2*], respectively. Each residue that differed from the germline gene was reverted back to the inferred germline residue. We then tested if the germline revertants of the antibodies in each group shared similar functional properties with their somatically-mutated counterparts. Each germline-revertant antibody was tested for binding to SARS-CoV-2 S6P_ecto_, SARS-CoV-2 RBD, or SARS-CoV-1 S2P_ecto_ proteins. The Group 1 germline revertant did not bind to SARS-CoV-2 S6P_ecto_ or SARS-CoV-1 S2P_ecto_. The Group 2 germline revertant maintained binding to both SARS-CoV-2 and SARS-CoV-1 proteins but exhibited lower binding avidity (higher EC_50_ values) than its matured counterparts COV2-2164 or CnC2t1p1_B10. The Group 3 germline revertant maintained binding to SARS-CoV-2 S6P_ecto_ and RBD proteins (**Fig. 4a, d**). Each germline revertant also bound to the surface of virus-infected cells (**Fig. 4b,d**). While none of the germline revertants exhibited neutralizing capacity (**Fig. 4c, d**), the Group 3 germline revertant showed a low level of ACE2 blocking (**Fig. 4d,e**).

**Figure 4.**
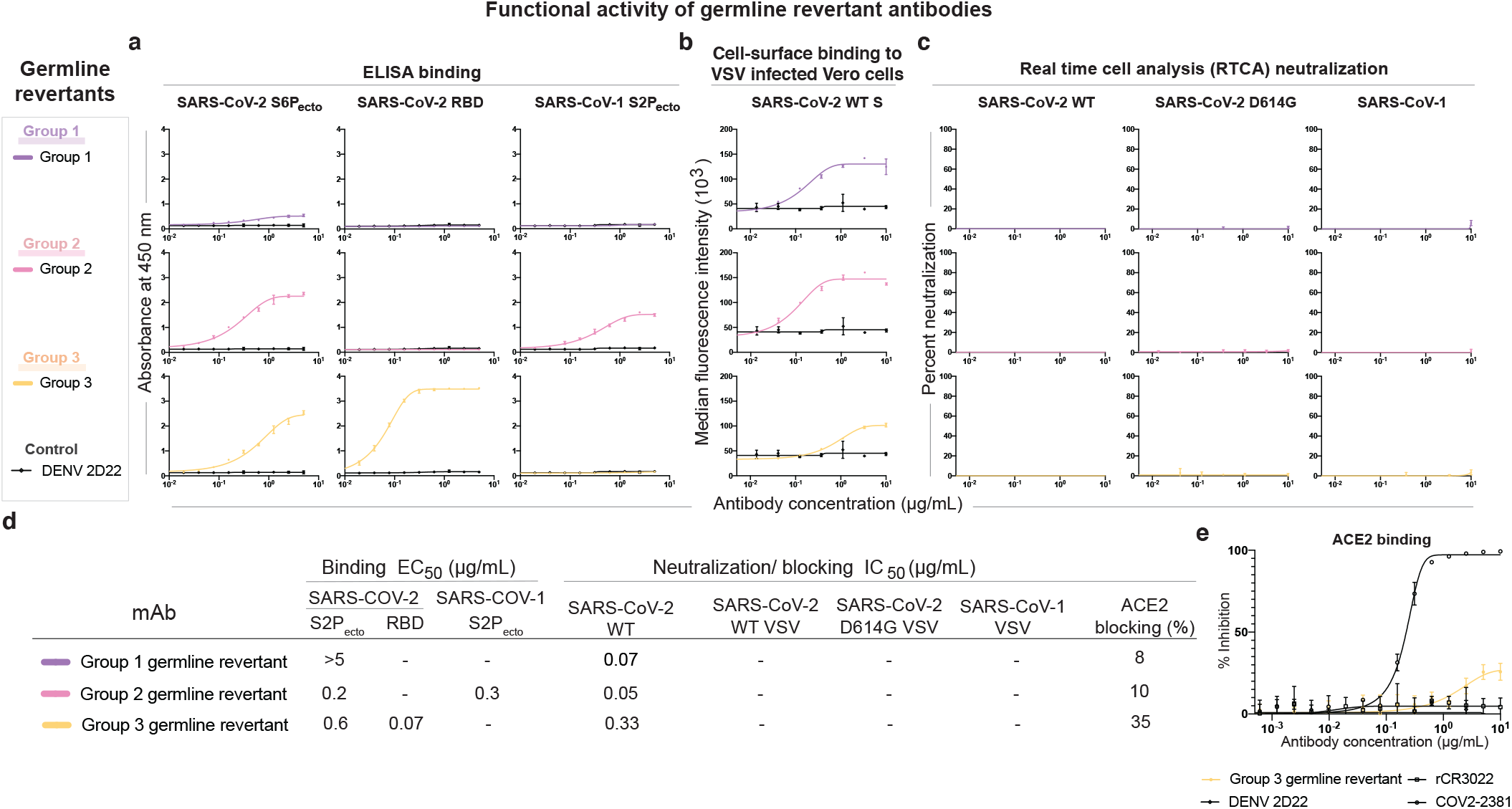
Germline-revertant antibody reactivity and functional activity. Group 1, 2, or 3 germline-revertant antibodies are shown in purple, pink, or yellow, respectively. DENV 2D22 was used as a control antibody for all assays, as shown in the lines in black. All experiments were performed in biological replicate and technical triplicate. Biological replicate from representative single experiment shown. a. Binding to SARS-CoV-2 S6P_ecto_, SARS-CoV-2 RBD, or SARS-CoV-1 S2P_ecto_ were measured by absorbance at 450 nm, as shown in the first three columns. b. Binding to Vero cells infected with VSV-SARS-CoV-2, measured by flow cytometric analysis and reported as median florescence intensity. c. Results for neutralization curves for replication-competent VSV chimeric viruses in real time cell analysis (RTCA) are shown in the next three columns, measured by percent neutralization calculated by normalized cell index. d. Binding EC_50_ and neutralization IC_50_ values for each of the assay curves in **Fig 5a**. All values are denoted as µg/mL. ACE2 blocking was determined by measuring amount of ACE2 with FLAG tag binding in the presence of each antibody, measured by binding of an anti-FLAG antibody. Percent blocking is shown, calculated by using ACE2 binding without antibody as 0% blocking. e. Inhibition binding curves for the Group 3 germline-revertant antibody. The starting antibody concentration used was 10 µg/mL and was titrated three-fold serially to obtain ACE2-blocking curves.

### COV2-2531 confers protection *in vivo*

MAbs can act by direct virus inactivation, but binding of some mAbs to the surface of virus-infected cells (**Fig. 2e, h**) suggested that these antibodies also might act through Fc-mediated functions. Therefore, it was important to test some public clonotypes *in vivo*. We tested the efficacy of these antibodies against SARS-CoV-2 *in vivo*. We used K18-hACE2 transgenic mice, which develop severe lung infection and disease after intranasal inoculation^48–50^. K18-hACE2 transgenic mice received either one antibody from Group 2 (COV2-2164), one antibody from Group 3 (COV2-2531), or an isotype-control antibody (DENV 2D22) via intraperitoneal injection (200 µg, 10 mg/kg) one day prior to intranasal inoculation with 10^3^ PFU of SARS-CoV-2 (WA1/2020). Mice treated with COV2-2531 were protected completely from weight loss (**Fig. 5a**) and showed reduced viral infection in the lung, nasal wash, heart, and brain (**Fig. 5b, c, d**) compared to the isotype-control antibody-treated group. However, mice treated with COV2-2164 were not protected from weight loss yet showed a reduction in viral load in the lung and brain (**Fig. 5b, e**) but not in the nasal wash and heart (**Fig 5c, d**). Thus, antibodies that compete for binding with the SARS-CoV-1 mAb rCR3022 can be elicited after SARS-CoV-2 infection, some of which can confer protection.

**Figure 5.**
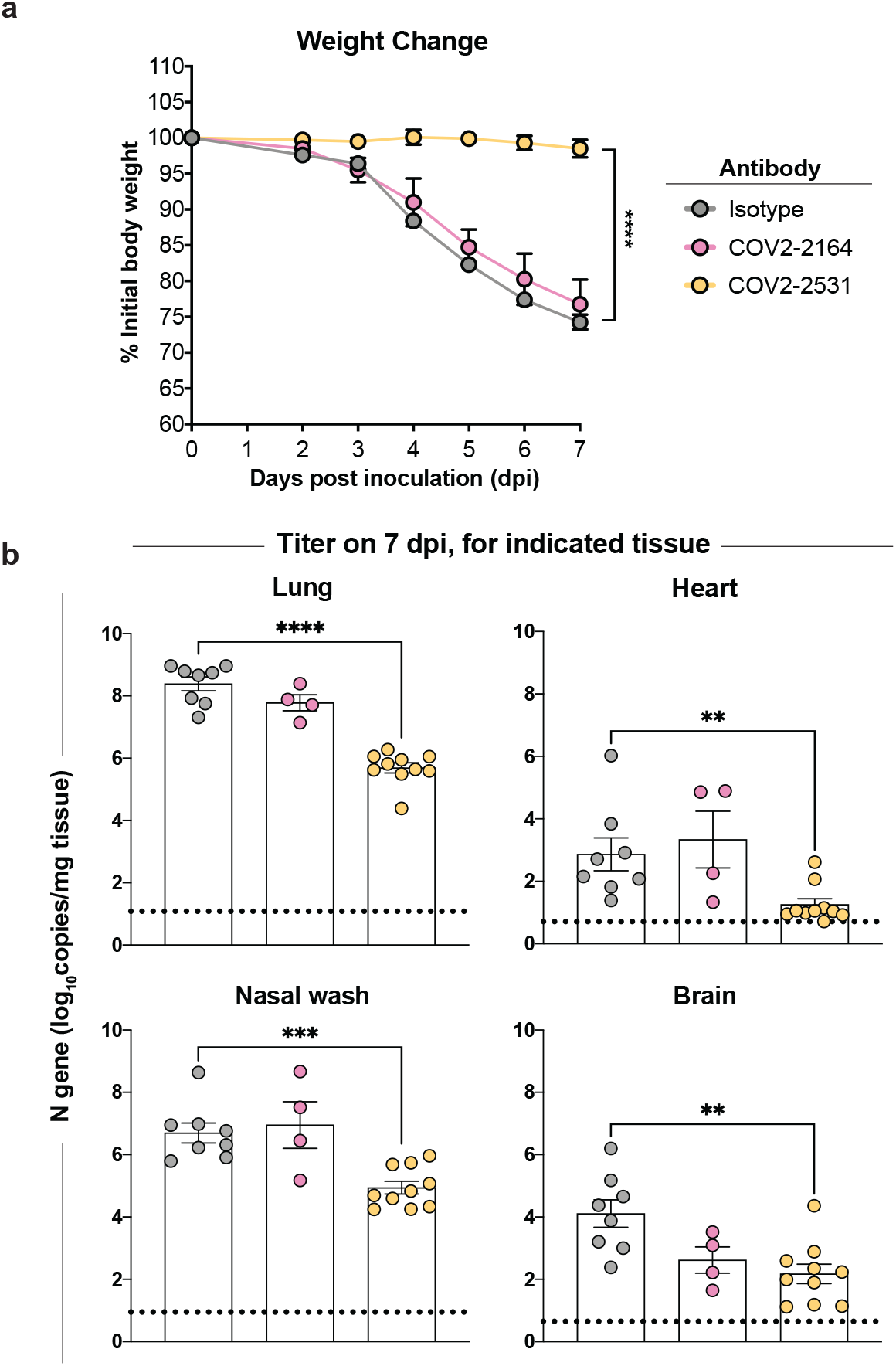
Antibody-mediated protection against SARS-CoV-2 challenge in mice. **a-e.** Eight-week-old male K18-hACE2 transgenic mice were inoculated by the intranasal route with 10^3^ PFU of SARS-CoV-2 (WA1/2020 strain). One day prior to infection, mice were given a single 200 μg dose of COV2-2351 or COV2-2164 by intraperitoneal injection. **a.** Weight change. Statistical analysis was performed only between isotype- and COV2-2351-treated groups. For isotype and COV2-2531 (mean ± SEM; n = 8-10, two experiments: unpaired t-test of area under the curve; **** *P* < 0.0001). For COV2-2164 (mean ± SEM; n = 8, two experiments) **b.** (**b-e**) Viral RNA levels at 7 days post-infection in the lung, nasal wash, heart, and brain as determined by qRT-PCR. For isotype and COV2-2531 (mean ± SEM; n = 8-10, two experiments: one-way ANOVA with Turkey’s post-test: ns not significant, * *P* < 0.05, *** *P* < 0.001, **** *P* < 0.0001, comparison to the isotype control mAb-treated group). For COV2-2164 (mean ± SEM; n = 8, two experiment).

### Public clonotypes shared between vaccine and convalescent responses to SARS-CoV-2 S protein

We hypothesized that SARS-CoV-2 mRNA vaccines might induce public clonotypes that are shared with those seen in convalescent individuals after natural infection. We obtained peripheral blood mononuclear cells from a volunteer 10 after first vaccine dose and 7 days after second vaccine dose with the Pfizer-BioNTech vaccine. Circulating plasmablasts were enriched directly from blood by negative selection using paramagnetic beads and purified further by flow cytometric sorting (**Fig. 6a, b**). Sorted plasmablasts were loaded on a Beacon microfluidics instrument for single-cell secreted antibody binding screening and antibody gene sequencing or in a Chromium single-cell microfluidics device (10X Genomics) followed by reverse transcription with PCR and sequence analysis to obtain paired antibody sequences. These antibody discovery workflows were described in detail previously^8^. Enzyme-linked immunospot (ELISpot) assay analysis revealed large increase in the frequency of S-reactive cells in the enriched plasmablast cell fraction on day 7 after the second vaccination compared to that on day 10 after the first vaccine dose, confirming induction of target-specific responses in this individual. SARS-CoV-2 S6P_ecto_-specific secreted antibodies were of IgG and IgA classes and accounted for >10% of total plasmablasts (**Fig. 6c**). Further, single-cell antibody secretion analysis of a total of 4,797 purified plasmablasts loaded on a Beacon microfluidics instrument (Berkeley Lights Inc.) revealed that a large fraction of SARS-CoV-2-reactive clones (included S6P_ecto_- and/or RBD-reactive clones) secreted RBD-specific IgG (**Fig. 6d**).

**Figure 6.**
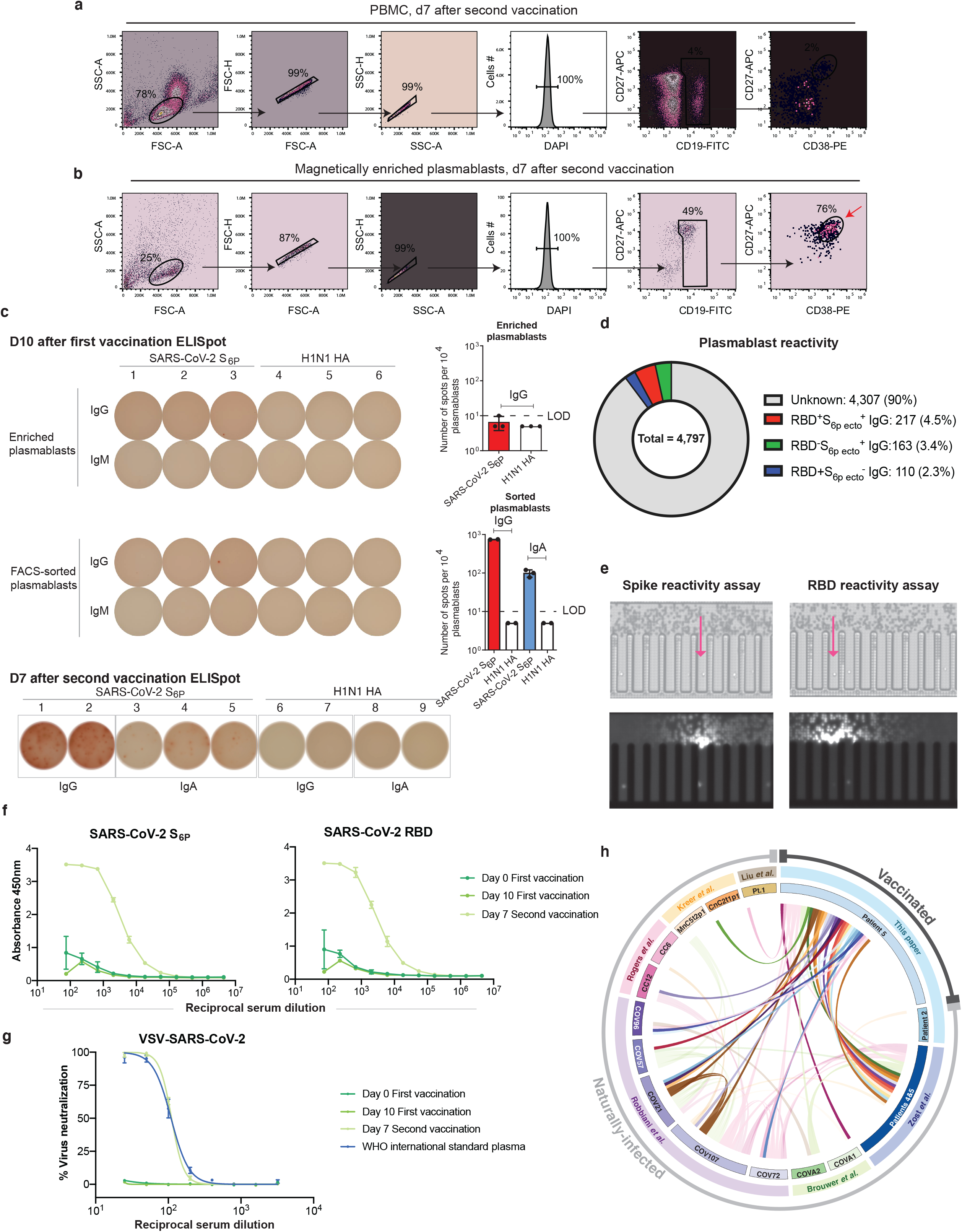
Analysis of vaccinated donor antibody response. a. Flow cytometric plots showing gating strategy to identify plasmablasts in total PBMC sample collected on day 7 after second vaccine dose (top panel) or identification of plasmablasts after direct enrichment from whole blood at the same time point using negative selection with paramagnetic beads (bottom panel). Blue arrow indicate enriched plasmablasts that were used for ELISpot analysis as in (b), and red arrow indicate plasmablasts (DAPI^-^CD19^lo^CD27^hi^CD38^hi^) that were FACS-sorted for single cell secretion and paired antibody sequencing studies. b. ELISpot analysis of SARS-CoV-2 S6P_ecto_-specific antibody secretion using enriched plasmablasts from blood collected on day 10 after the first vaccine dose (IgG), and day 7 after the second vaccine dose (IgG and IgA). A/Darwin/42/2020 H1N1 influenza virus hemagglutinin (HA) was used as a control for specificity of the plasmablast response. Wells with spots (left) and number of SARS-CoV-2 S6P_ecto_-specific responses detected (right) is shown. Dotted line indicates values below limit of the detection (LOD=10 spots per 10^4^ cells), that were set up to 5 spots per 10^4^ cells. c. FACS-sorted plasmablasts were loaded on a Beacon optofluidic instrument and assessed for binding to S6P_ecto_ or RBD-coated beads using single cell antibody secretion analysis. Bright field images of the Beacon instrument chip with individual plasmablasts loaded into the pens of the chip are shown for the selected fields of view for each screening condition. False-color fluorescent images from the same fields of view showing binding of the detection anti-human Alexa-Fluor-568-labeled antibody to the S6P_ecto_ or RBD-coated beads that captured human antibodies secreted by single plasmablasts (visualized as a plume from the beads that loaded into the channel of the chip). d. Pie chart representation showing frequency of RBD and SARS-COV-2 S6P_ecto_ reactive (red), SARS-COV-2 S6P_ecto_ reactive only (green), or RBD reactive only (blue) plasmablasts identified as in c. Fraction of cells that did not react to either SARS-COV-2 RBD or S6P_ecto_ is shown in grey. e. Flow-cytometry-sorted plasmablasts were loaded on a Beacon instrument and assessed for binding to S6P_ecto_ or RBD-coated beads using single-cell antibody secretion analysis. Bright field images of the Beacon instrument chip with individual plasmablasts loaded into the pens of the chip are shown for the selected fields of view for each screening condition (top). False-color fluorescent images from the same fields of view (bottom) showing binding of the detection anti-human Alexa-Fluor-568-labeled antibody to the S6P_ecto_ or RBD-coated beads that captured human antibodies secreted by single plasmablasts (visualized as a plume from the beads that loaded into the channel of the chip). Arrow indicates cells secreting antigen-reactive IgG antibodies f. ELISA binding to SARS-CoV-2 S6P_ecto_, of serum from patient 5 at days 0 of first vaccination, day 10 after first vaccination, day 7 after second vaccination, or day 28 after second vaccination were measured by absorbance at 450 nm. Serum was diluted 1:75 and then diluted serially three-fold. Experiment performed in biological replicate and technical triplicate. Biological replicate from representative single experiment shown. g. Neutralization curves of serum from patient 5 at days 0 of first vaccination, day 10 after first vaccination, or day 7 after second vaccination. A WHO International standard for anti-SARS-CoV-2 human immunoglobulin was used as the positive control. Serum was diluted starting at a 1:25 dilution, then diluted serially two-fold. Experiment performed in technical triplicate. h. Circos plot indicating public clonotypes identified in this paper. The more opaque ribbons within the circle represent public clonotypes that are shared between the vaccinated donor and convalescent donors after natural infection. Translucent ribbons indicate public clonotypes shared between convalescent infection individuals. The individuals from whom sequences were derived are indicated on the inner circle. The published sources from which the sequences were obtained are shown on the second circle. The outside circle indicates whether the individuals were naturally-infected or vaccinated.

We also analyzed antibody reactivity and neutralization in serum collected on the day before vaccination (day 0), on day 10 after the first vaccine dose, on day 7 after the second vaccine dose, and on day 28 after the second vaccine dose. The reactivity of serum antibodies to both SARS-CoV-2 S6P_ecto_ and SARS-CoV-2 RBD was measured by ELISA for binding (**Fig. 6f**) and by VSV-SARS-CoV-2 neutralizing assay (**Fig. 6g**). Binding and neutralizing activities steadily increased over time, with maximal activity detected on day 28 after the second vaccine dose.

From single-cell antibody variable gene sequencing analysis, we obtained 725 paired heavy and light chain sequences from plasmablasts following primary immunization and 8,298 paired sequences from plasmablasts following the second dose of vaccine. The same procedure was carried out on a sample collected 35 days after onset of symptoms from a convalescent individual with confirmed SARS-CoV-2 infection. This individual’s serum had been determined previously to contain neutralizing antibodies^8^. Single-cell antibody secretion analysis revealed that a minor fraction (0.5%) of total plasmablasts produced S-protein-reactive antibodies. We identified 1,883 paired heavy and light chain antibody sequences for this specimen.

Antibody sequences identified in these new studies and sequences we collected from previous SARS-CoV-2 antibody discovery studies were clustered as described in **Fig. 1**. We identified a total of 37 public clonotypes, 26 of which represented clonotypes shared between antibodies isolated from the vaccinee and individuals with exposure history to natural SARS-CoV-2 infection (**Fig. 6h**). The antigen-binding specificity of each group was inferred through review of data in each respective publication in which the antibodies were reported. We determined that 14 of the 26 newly-identified shared clonotypes encoded antibodies specific to the SARS-CoV-2 S protein. Within that panel of mAbs, 8 of 26 clonotypes reacted with SARS-CoV-2 RBD protein, and 6 of the 26 public clonotypes cross-reacted with both SARS-CoV-1 and SARS-CoV-2 (**Fig. 7)**. Most antibodies shared in public clonotypes were IgG, with a subset of IgAs noted. This finding shows that the Pfizer-BioNTech vaccine induces many antibodies that are genetically similar to ones elicited through natural SARS-CoV-2 infection, including multiple public clonotypes in convalescent donors encoded by commonly used V_H_ genes such as *IGHV3-53*, *IGHV3-66*, *IGHV1-58*, *IGHV3-30*, and *IGHV3-30-3*. Additionally, of the 37 total public clonotypes, 16 bound to RBD, and of these, 11 of 16 were neutralizing. All neutralizing public clonotypes recognized RBD. However, of the 37 public clonotypes identified, 21 are directed to antigenic sites other than the RBD, including ones described here directed to the S2 domain. It is likely that although a substantial portion of neutralizing public clonotypes is directed to the RBD, non-RBD-targeted and non-neutralizing public clonotypes may make up an even larger portion of an individual’s response to either vaccination or infection. Overall, these results suggest that many of the public clonotypes observed in previously infected individuals likely are found in vaccinated individuals.

**Figure 7.**
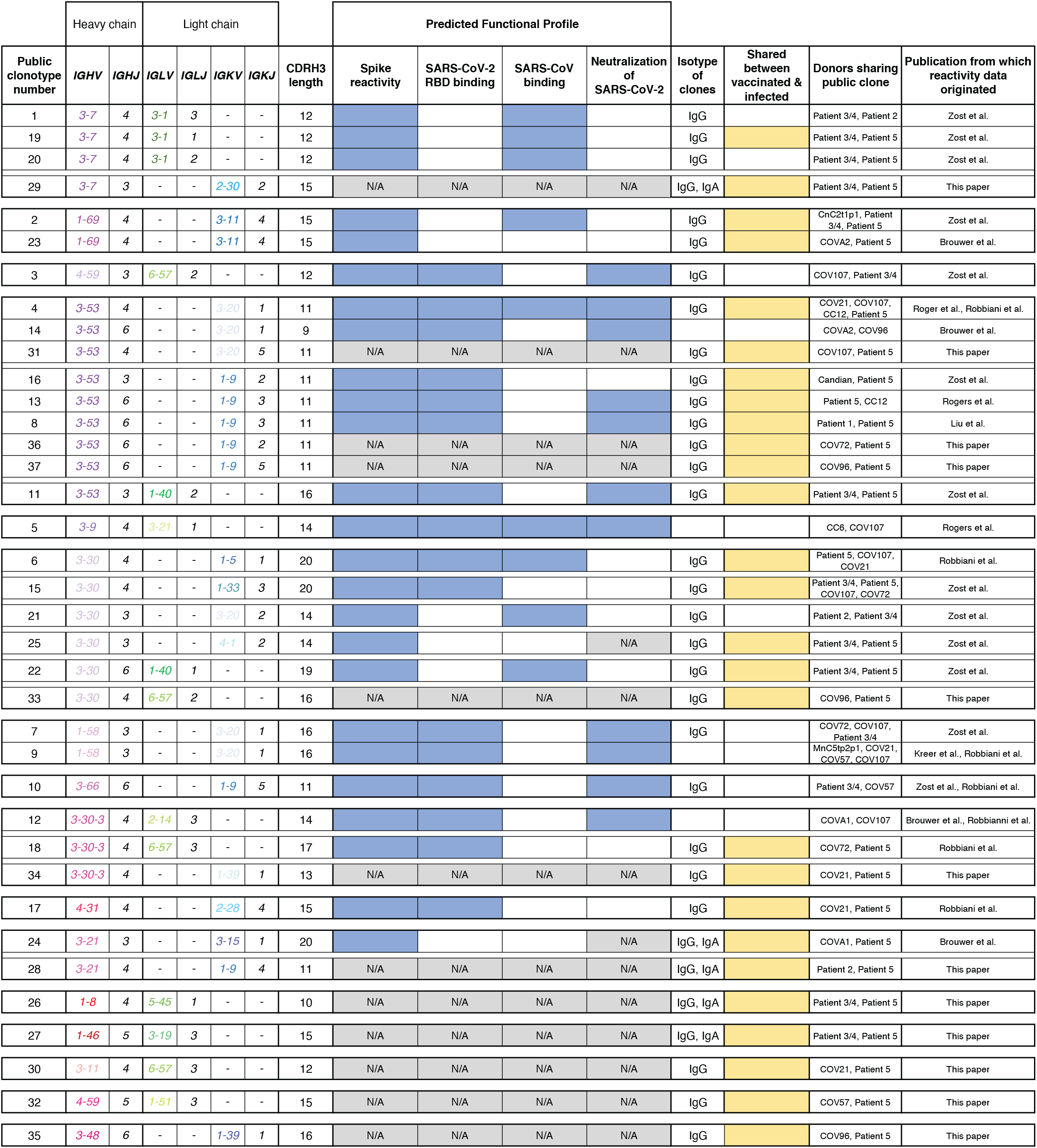
Identification of public clonotypes shared between naturally-infected individuals and a vaccinated donor. a. Table showing all public clonotypes identified. Gene usage for each clone or CDRH3 length are shown in columns 2 or 3. Reactivity profiles obtained from published sources are shown for comparative purposes. Blue indicates positive reactivity, while white indicates that binding reactivity or neutralization was not detected. Grey indicates reactivity profile was not found in either publication and therefore is unknown. Isotypes of antibodies in each group are listed in the eighth column. If the group contained sequences from both vaccinated and infected individuals, it was denoted in yellow. White was used for clonotypes that were shared only between convalescent individuals following natural infection.

## DISCUSSION

The high number of identified public B cell clonotypes in the response to SARS-CoV-2 infection or vaccination is striking, and the frequency of public clonotypes identified here is much higher than in randomly sampled B cells in a convalescent donor^8^. Many public clonotypes are shared between both infected and vaccinated individuals. Public clonotypes appear to be induced by each of the currently known antigenic sites on the S protein and are found in both the neutralizing and non-neutralizing repertoires. Some clonotypes in the shared SARS-CoV-2 response appear preconfigured in the germline state to recognize particular S epitopes, and this recognition likely is driven by particular structural features on S. The scale of available relatively large repertoire data for SARS-CoV-2 enabled us to identify many public clonotypes. The SARS-CoV-2 pandemic has resulted in comprehensive studies of antibody responses to SARS-CoV-2 S protein, with many groups identifying large panels of mAbs, including potently neutralizing ones^8–12,38,40,41^. These discovery efforts have led to the identification of large paired heavy and light chain antibody variable gene sequence data sets for B cells specific to SARS-CoV-2, and the data has been made public at a scale unlike that for any other virus. In this study we compared the sequences of more than 14,000 paired B cell sequences encoding antibodies to S protein of SARS-CoV-2. Likely, this influx in the availability of paired antibody gene sequences from a multitude of donors contributed to our ability to identify an unexpectedly high number of paired-sequence public clonotypes. It will be interesting in future to use paired sequencing to determine if the scale of shared repertoire we observed here for SARS-CoV-2 is a more pronounced feature of the response to this particular virus than that of other viruses that have been studied for shared clones, such as HIV-1, influenza, and hepatitis C. Previous studies have identified public clonotypes in the response to these other viral pathogens, for example a recent study of response to HIV^26–29^ in which 27 public clonotypes were described with unpaired sequencing using only the heavy chain CDR3 sequence and V_H_/J_H_ gene usage^26^.

Several neutralizing public clonotypes have be identified previously, most commonly clonotypes encoded by the closely related heavy chain genes *IGHV3-53*, I*GHV3-66*^32,37^, *IGHV1-2*^*51*^*, and IGHV3-30*^9^. Structural features of these public clonotypes likely drive the frequent selection of such clones, such as the canonical configuration of aromatic residues in the public clonotype *IGHV1-58 + IGHJ3* and *IGKV3-20 + IGKJ1* that engages the SARS-CoV-2 RBD F486 residue^33^. Members of this public clonotype, such as COV2-2196, engage the RBD using predominantly germline-encoded residues in both the heavy and light chain^*9,33,34,39,52*^. Identification of public clonotypes from multiple donors suggest these antibodies could contribute to humoral responses that mediate protection if they appear not only in memory B cells but also as antibodies from plasma cells secreted into the serum^53^. The high prevalence of public clonotypes elicited to the SARS-CoV-2 S trimer may contribute to the high efficacy of S-encoding mRNA vaccines in large populations.

The recognition pattern of public clonotypes may predict the emergence of particular antibody escape virus variants. If diverse individuals independently make the same antibody in response to an antigen, there could be a constant and collective selective pressure on that epitope, resulting in a high potential for escape variants at that site. For example, while *IGHV3-53-* and *IGHV3-66*-encoded public clonotypes have been described in numerous individuals, neutralization of these antibodies is impacted adversely by the K417N or K417T substitutions present in the B.1.351 or P.1 SARS-CoV-2 variants of concern, respectively^54^. A similar case was described for *IGHV1-2-*encoded antibodies that target the RBD and *IGHV1-24*-encoded antibodies that target the NTD. These antibodies are found in the serum of convalescent individuals^53^, but neutralization of these antibodies is negatively affected for 501Y.V2 variant viruses^55^. A possible explanation for the selective pressure that led to the emergence and propagation of these variants is the humoral immunity mediated by these public clonotypes.

The new Group 3 public clonotype neutralizing and protective antibodies described here bind to the cryptic face of the RBD and compete with the SARS-CoV-2 non-neutralizing mAb CR3022. Neutralizing antibodies that bind to the more conserved base of the RBD are of interest, as these sites are largely unaffected by common mutations in the variants of concern such as E484K, K417N, and N501Y^54^. Importantly, recent work has identified a B.1.1.7 variant with a deletion of RBD residues 375-377. This deletion disrupts the epitope of CR3022, yet appears to be functionally tolerated^56^. As Group 3 antibodies share a similar epitope, with critical residues of COV2-2531 and C126 being K378 and A372, but with additional critical residues of Y369, N370, F374, and P384 identified for C126, this deletion might abrogate binding of antibodies from this public clonotype. Further study of public clonotypes may give insight into the most likely sites of future major antigenic changes in circulating field strains.

While public clonotypes have been described that recognize the RBD^33,9,32,37,51^ and NTD^53,57,58^ of the S trimer, to our knowledge, those specific to the S2 domain have not been described. In this study, we identified two public clonotypes, designated here as Groups 1 and 2, which target the S2 domain of the S trimer. These mAbs do not neutralize, but they react with S proteins of both SARS-CoV-2 and SARS-CoV-1. It is likely that these S2 epitopes are the target of non-neutralizing antibodies in multiple individuals following infection or vaccination. Previous studies have identified broadly immunogenic epitopes that are conserved in the functional domains of the SARS-CoV-2 S trimer S2 domain, including cross-reactivity to endemic coronaviruses, and therefore these findings have important implications for antibody and vaccine design^44^. Rational reverse vaccinology approaches such as structure-based design of targeted antibody epitopes offer an opportunity to elicit or prevent boosting of neutralizing or non-neutralizing antibodies as desired^59^. The S2 region of the S trimer may be more capable of recruiting preexisting memory B cells for diverse coronaviruses, since the S2 domain is more conserved for functionally important sites such as the heptad repeat regions and fusion loop^60^. With a variety of public clonotype reactivities occurring to regions other than the RBD, it is likely that there are many additional public clonotypes that recognize the S2 domain or other regions of the S trimer. Although the S2-reactive public clonotypes described here (Groups 1 and 2) did not neutralize or protect, future studies should investigate if additional non-RBD targeted public clonotypes can induce protection.

We propose that there are essentially four classes of public clonotypes (**Extended data Fig. S8**): (1) neutralizing public clonotypes that bind to relatively invariant sites on S, (2) neutralizing public clonotypes that bind to sites that tolerate high sequence variability, (3) non-neutralizing public clonotypes that target relatively invariant sites, and (4) non-neutralizing public antibodies that target variable sites. The first class of antibodies is likely the most protective class in a population, as these mAbs neutralize and recognize residues unlikely to be sustained with mutations due to loss of viral fitness. An example of this class would be *IGHV1-58-*encoded antibodies as described previously^33^. Many public clones currently identified for SARS-CoV-2 are categorized in the second class. While these clones initially offer protection, this property could be lost as widespread selective pressure on the virus is exerted on a region with genetic and structural plasticity. Examples of this group were discussed here, such as *IGHV3-53-* and *IGHV3-66*-encoded antibodies that target the RBD^54^. Here, we described three new public clonotypes following natural infection (Groups 1, 2, and 3) and a total of 29 new clonotypes after mRNA vaccination. Public clonotype Groups 1 and 2 fall into the third class of antibodies described here (non-neutralizing antibodies that target invariant sites), and public clonotype Group 3 antibodies falls into the second class (neutralizing public clonotypes that bind to variable sites). Future public clonotypes to SARS-CoV-2 could be binned with this four-quadrant scheme to better understand how public clonotypes contribute to humoral immunity against COVID-19.

Understanding the antibody response that is shared between convalescent and vaccinated individuals also will be of continued interest as the percentage of vaccinated individuals increases in the facing of emergence of new viral variants of concern. The understanding of viral epitopes that induce protective antibodies in multiple individuals has implications for predicting the most common responses to new vaccines in large populations. The emergence of SARS-CoV-2 variants with acquired mutations in epitopes for neutralizing antibodies, including antibody regimens currently authorized for EUA, is a cause for concern^61–64^. Our analyses of public clonotypes after natural infection and vaccination and their shared epitope targets may predict sites of future major antigenic changes in the S trimer.

## Data and materials availability

Sequence Read Archive deposition for the public clonotypes identified is deposited at the NCBI: PRJNA511481. All other data are available in the main text or the supplementary materials. Requests for reagents may be directed to and be fulfilled by the Lead Contact: Dr. James E. Crowe, Jr. Materials reported in this study will be made available but may require execution of a Materials Transfer Agreement.

## Software availability

The computational pipeline for the clustering analysis as well as the script to create heatmaps is available on GitHub: https://github.com/eccelaine/COV2-Public-Clonotypes. PyIR script used to determine sequence characteristics of each antibody is available here: https://github.com/crowelab/PyIR.

## Acknowledgments

Electron microscopy data collection was collected at the Center for Structural Biology Cryo-EM facility at Vanderbilt University. We thank Cinque Soto for technical advice on study of public clonotypes. We thank Rachel Nargi, Joseph Reidy, Erica Armstrong, and Christopher Gainza for support in antibody purifications. We also thank Berkeley Lights and Jonathan Didier for expert technical support, and STEMCELL Technologies and Aida Mayhew for supplying crucial B cell enrichment reagents. We thank Jem Uhrlaub, Brendan Larson, and Dr. Mike Worobey (University of Arizona) for preparation and sequencing of early-passage SARS-CoV-2 stocks. We thank Eun-Hyung Lee, Doan C. Nguyen, and Ignacio Sanz from Emory University for sharing plasmablast survival medium that promotes antibody secretion. We thank the anonymous donors of the plasma samples for their consent which has allowed WHO International standard for anti-SARS-CoV-2 human immunoglobulin to be prepared; we express our gratitude to those who have coordinated the collection of the convalescent plasma: Malcom Semple (University of Liverpool, UK), Lance Turtle (University of Liverpool, UK), Peter Openshaw (Imperial College London, UK) and Kenneth Baillie (University of Edinburgh) on behalf of the ISARIC4C Investigators; Heli Harvala Simmonds and David Roberts (National Health Service Blood and Transplant, UK). We also thank NIBSC Standards Production and Development staff for the formulation and distribution of materials.

## Author contributions

Conceptualization, E.C.C. and J.E.C.; Investigation, E.C.C, P.G., S.J.Z., N.S., M.S., C.S., E.S.W., C.R.C., C.A.T., S.L., S.K.C., E.B., R.S.N., J.R., E.A., C.G., S.D., ., A.T., J.R., R.E.S., E.D.; Writing first draft: E.C.C. and J.E.C; All authors edited the manuscript and approved the final submission; Supervision, B.J.D., R.H.C., C.T., M.S.D., J.E.C.; Funding acquisition, M.S.D., J.E.C.

## Additional information

**Supplementary information** is available for this paper.

**Correspondence and requests for materials** should be addressed to J.E.C.

---

**Extended Data Fig. 1. Clustering schematic to identify public clonotypes.** Schematic of how sequences were binned and clustered to identify public clonotypes.

**Extended Data Fig. 2. Controls for ELISA and neutralization assays.**

**a.** Positive or negative controls used for testing antibody binding in ELISA to SARS-CoV2-S6P_ecto_, SARS-CoV-1 S2P_ecto_, or SARS-CoV-2 RBD proteins. The positive control antibody COV2-2381 binds to SARS-CoV-2 S2P_ecto_ and RBD but not to SARS-CoV-1 S2P_ecto_, and the positive control antibody rCR3022 also binds to SARS-CoV-1 S2P_ecto_.

**b.** Positive or negative controls used for replication-competent chimeric VSV neutralization assays. COV2-2381 was used as a positive control for SARS-CoV-2 WT and D614G, whereas rCR3022 was used as a positive control for SARS-CoV-1.

**Extended Data Fig. 3. Staining of dsRNA intensity.** Staining of dsRNA intensity split into DAPI stain, dsRNA stain, and the two merged for each antibody group.

a. Staining for Group 1 antibodies.

b. Staining for Group 2 antibodies.

c. Staining for Group 3 antibodies.

d. Staining for control antibodies. 2D22^46^ is used as a negative control antibody. COV2-2130^42^ is used as a positive control antibody.

**Extended Data Fig. 4. Negative stain complexes of each public clonotype**

a. Negative stain EM of SARS-CoV-2 S6P_ecto_ protein in complex with Fab forms of COV2-2002 or COV2-2333.

b. Negative stain EM of SARS-CoV-2 S6P_ecto_ protein in complex with Fab forms of COV2-2164 or CnC2t1p1_B10.

c. Negative stain EM of SARS-CoV-2 S6P_ecto_ protein in complex with Fab forms of COV2-2531 or C126.

**Extended Data Fig. 5. Control reagents for detection of antibody binding to membrane-anchored S protein in cell-surface antigen-display assays,**

a. Gating strategy used for cell-surface antigen-display experiment. The first gate is for all cells, the second gate is for infected cells, and the third gate is for antibody binding to infected cells.

b. Controls used for cell-surface antigen-display antibody binding experiment. Cell-only control in shown in light grey. The unrelated mAb DENV 2D22 was used as an antibody negative control, shown in dark grey. The mAb COV2-2381 shown in dark blue and mAb rCR3022 shown in turquoise were used as positive antibody controls.

c. Histogram of data obtained using infected or uninfected cells. Infected cells are shown in light grey, and uninfected cells are shown in dark grey.

d. Group 1, 2, or 3 antibody binding to infected cells. The antibody concentration used was 10 µg/mL for all antibodies.

e. Group 1, 2, or 3 germline-revertant antibody binding to infected cells. The antibody concentration used was 10 µg/mL for all antibodies.

**Extended Data Fig. 6. Primary data for alanine mutagenesis screening.** Binding values for mAbs on the SARS-CoV-2 S protein alanine scan library. The binding values at critical mutant clones for

a. Group 1 (COV2-2002 in light purple and COV2-2333 in dark purple) and Group 2 (COV2-2164 in pink and CnC2t1p1_B10 in red)antibodies are shown as a percentage of mAb binding to wild-type (WT) SARS-CoV-2 spike protein and are plotted with the range (highest-minus lowest binding value) of at least two measurements.

b. Group 3 (COV2-2531 in light orange and C126 in dark orange) antibodies are shown as a percentage of mAb binding to wild-type (WT) SARS-CoV-2 spike protein and are plotted with the range (highest-minus lowest binding value) of at least two measurements.

**Extended Data Fig. 7. Overlay of CR3022 structure with Group 3 antibodies when bound to RBD**

a. The structures for the RBD domains for both SARS-CoV-2 and SARS-CoV-1 were overlaid. The epitope of rCR3022 is highlighted in orange (from Yuan *et al*.). Light orange dots denote the binding residues for mAb COV2-2531, and dark orange dots denote the binding residues for C126. Figure adapted from previous study^47^.

**Extended Data Fig. 8. Proposed classes of public clonotypes to SARS-CoV-2.** There are four proposed classes of public clonotypes to SARS-CoV-2, separated by the relationship between variability of targeted epitope (y-axis) and the neutralizing potency (x-axis) of each antibody clonotype.

## Materials and Methods

### Research participants

We studied three patients (patient 2, 3 and 4) with in North America with laboratory-confirmed symptomatic SARS-CoV-2 infections that we have described previously^8^. We studied one patient (a 59-year-old male) who received Pfizer-BioNTech vaccine. The studies were approved by the Institutional Review Board of Vanderbilt University Medical Center.

### Cell lines

Vero E6 (ATCC, CRL-1586) cells were maintained at 37°C in 5% CO_2_ in Dulbecco’s minimal essential medium (DMEM) containing 10% heat inactivated fetal bovine serum (FBS), 10 mM HEPES pH 73, 1 mM sodium pyruvate, 1× non-essential amino acids, and 100 U/mL of penicillin-streptomycin. ExpiCHO cells (Thermo Fisher Scientific, A29127) were maintained at 37°C in 8% CO_2_ in ExpiCHO Expression Medium (Thermo Fisher Scientific, A2910002). Mycoplasma testing of cell lines was performed on monthly basis using a PCR-based mycoplasma detection kit (ATCC, 30-1012K), with negative results at each testing. Calu-3 (ATCC, HTB-55) cells were maintained at 37°C in 5% CO_2_ in DMEM with high glucose and L-glutamine (Gibco 11965092), containing 10% heat inactivated fetal bovine serum (FBS), and 100 U/mL of penicillin-streptomycin.

### Viruses

The generation of a replication-competent VSV expressing SARS-CoV-2 S protein with a 21 amino acid C-terminal deletion that replaces the VSV G protein (VSV-SARS-CoV-2) was described previously^65^. The S protein-expressing VSV virus was propagated in MA104 cell culture monolayers (African green monkey, ATCC CRL-2378.1)^65^. Viral stocks were titrated on Vero E6 cell monolayer cultures by visualizing VSV plaques using neutral red staining. VSV-SARS-CoV-2/D614G was introduced by site directed mutagenesis. The 2019n-CoV/USA_WA1/2019 isolate of SARS-CoV-2 was obtained from the US Centers for Disease Control (CDC). Infectious stocks were propagated by inoculating Vero CCL81 cells. Supernatant was aliquoted and stored at -80°C. The University of Arizona group obtained the USA-WA1/2020 isolate of SARS-CoV-2 from WRCEVA. Early passage virus stock was generated by a single passage on Vero CCL81 for 48 h. Infected cell lysate and culture supernatant was combined, subjected to one freeze-thaw, and then centrifuged to pellet cell debris. The stock was titered to ∼3 x10^6^ PFU/mL by standard plaque assay on Vero CCL81 cells. Nanopore sequencing of these early passages confirmed the genome sequence was identical to the Genbank WA1/2020 sequence (MN985325.1), with no mutations in the spike furin cleavage site. All work with infectious SARS-CoV-2 was performed in Institutional Biosafety Committee-approved BSL3 or A-BSL3 facilities at Washington University School of Medicine or University of Arizona, using appropriate positive pressure air respirators and protective equipment.

### Clustering for identification of public clonotypes

Publicly available paired sequence sets of antibody genes were obtained^8,10–12,38–40^. Together with sequences derived from this paper, were first binned all sequences by the same heavy chain V and J genes. Following sequences then were clustered according to 70% sequence similarity on their CDRH3 nucleotide sequence. Lastly, sequences then were binned together again if they used the same light chain V and J genes. Clusters of sequences containing sequences from two or more donors were determined to be public clonotypes.

### Heat map generation

All sequences that were identified to be public clonotypes were analyzed with PyIR^66^ to identify the V and J genes. The number of sequences with corresponding V and J genes on the heavy and light chains were counted. These frequency counts then were plotted onto the heatmap using Python Seaborn Library.

### Antibody production and purification

Sequences of mAbs were synthesized using a rapid high-throughput cDNA synthesis platform (Twist Bioscience) and subsequently cloned into an IgG1 monocistronic expression vector (designated as pTwist-mCis_G1) for mAb secretion from mammalian cell culture. This vector contains an enhanced 2A sequence and GSG linker that allows simultaneous expression of mAb heavy- and light-chain genes from a single construct upon transfection^67^. We performed transfections of ExpiCHO cell cultures using the Gibco ExpiCHO Expression System and protocol for 50mL mini bioreactor tubes (Corning) as described by the vendor. Culture supernatants were purified using HiTrap MabSelect SuRe (Cytiva, formerly GE Healthcare Life Sciences) on a 24-column parallel protein chromatography system (Protein Biosolutions). Purified monoclonal antibodies were buffer exchanged into PBS, concentrated using Amicon Ultra-4 50-kDa centrifugal filter units (Millipore Sigma) and stored at 4°c until use.

### Expression and purification of recombinant receptor binding domain (RBD) of SARS-CoV-2 S protein

For electron microscopy imaging of S protein in complex with Fab forms of human mAbs, we expressed a variant of S6P_ecto_ protein containing a C-terminal Twin-Strep-tag, similar to that described previously^8^. Expressed protein was incubated with BioLock (IBA Lifesciences) and then isolated by Strep-tag affinity chromatography on StrepTrap HP columns (GE Healthcare), followed by size-exclusion chromatography on TSKgel G4000SWXL (TOSOH) if needed.

### ELISA binding assays

Wells of 384-well microtiter plates were coated with purified recombinant SARS-CoV-2 S6P_ecto_, SARS-CoV-2 RBD, or SARS-CoV S2P_ecto_ at 4°C overnight. Plates were blocked with 2% non-fat dry milk and 2% normal goat serum in DPBS containing 0.05% Tween-20 for 1 h. All antibodies were diluted to a concentration of either 0.4 µg/mL for the matured antibodies or 5 µg/mL for the germline-revertant antibodies. Antibodies were diluted in two-fold dilutions until binding was no longer detected. Bound antibodies were detected using goat anti-human IgG conjugated with horseradish peroxidase and TMB substrate. The reaction was quenched with 1N hydrochloric acid once color was developed. The absorbance was measured at 450 nm using a spectrophotometer (Biotek).

### Cell-surface antigen-display assay

Vero cell monolayers were monitored until 80% confluent and then inoculated with VSV-SARS-CoV-2 V (WA1/2020 strain) at an MOI of 0.5 in culture medium (DMEM with 2% FBS). For a T-225 flask, 10 mL of diluted VSV-SARS-CoV-2 virus was added to the monolayer, then incubated for 40 min. During the incubation, the flask was gently rocked back and forth every 10 min to ensure even infection. Following, the incubation the flask volume was topped off to 30 mL with 2% FBS containing DMEM and incubated for 14 h. Cells were monitored for CPE under a microscope, were trypsinized and washed in fluorescence activated cell sorting (FACS) buffer. 100,000 infected cells were seeded per well to stain with respective antibodies. All antibody was diluted to 10 µg/mL in FACS buffer, and then serially diluted 3-fold 7 times to stain for antibodies that react to cell-surface-displayed S protein. Infected cells then were resuspended in 50 µL of diluted antibody. Antibody binding was detected with anti-IgG Alexa-Fluor-647-labelled secondary antibodies. Cells were analyzed on an iQue cytometer for staining first by gating to identify infected cells as indicated by GFP-positive cells, and then gated for secondary antibody binding.

### Real-time cell analysis (RTCA) neutralization assay

To determine neutralizing activity of purified antibodies or human serum, we used real-time cell analysis (RTCA) assay on an xCELLigence RTCA MP Analyzer (ACEA Biosciences Inc.) that measures virus-induced cytopathic effect (CPE)^8,43,58^. Briefly, 50 μL of cell culture medium (DMEM supplemented with 2% FBS) was added to each well of a 96-well E-plate to obtain background reading. A suspension of 18,000 Vero cells in 50 μL of cell culture medium was seeded in each well, and the plate was placed on the analyzer. Measurements were taken automatically every 15 min, and the sensograms were visualized using RTCA software version 2.1.0 (ACEA Biosciences Inc). SARS-CoV-2 S VSV, SARS-CoV-2 S D614G VSV, or SARS-CoV-1 (∼0.02 MOI, ∼120 PFU per well) was mixed 1:1 with a respective dilution of mAb or heat-inactivated human serum in a total volume of 100 μL using DMEM supplemented with 2% FBS as a diluent and incubated for 1 h at 37°C in 5% CO_2_. At 16 h after seeding the cells, the virus-mAb mixtures were added in replicates to the cells in 96-well E-plates. Triplicate wells containing virus only (maximal CPE in the absence of mAb) and wells containing only Vero cells in medium (no-CPE wells) were included as controls. Plates were measured continuously (every 15 min) for 48 h to assess virus neutralization. Normalized cellular index (CI) values at the endpoint (48 h after incubation with the virus) were determined using the RTCA software version 2.1.0 (ACEA Biosciences Inc.). Results are expressed as percent neutralization in a presence of respective mAb relative to control wells with no CPE minus CI values from control wells with maximum CPE. RTCA IC_50_ values were determined by nonlinear regression analysis using Prism software.

### Competition-binding ELISA

Wells of 384-well microtiter plates were coated with purified recombinant SARS-CoV-2 S6P_ecto_ protein at 4°C overnight. Plates were blocked with 2% bovine serum albumin (BSA) in DPBS containing 0.05% Tween-20 for 1 h. Each antibody was diluted to a concentration of 10 µg/mL. Next, biotinylated antibodies were diluted to 2.5 µg/mL and added to the primary antibody solution without washing the plate to a final concentration of 0.5µg/mL. Biotinylated antibody binding was detected with horseradish peroxidase-conjugated avidin (Sigma) and developed with TMB. The reaction was quenched with 1N hydrochloric acid once color was developed. Absorbance was measured at 450 nm using a spectrophotometer.

### ACE2 blocking assay

Wells of 384-well microtiter plates were coated with purified recombinant SARS-CoV-2 S6P_ecto_ protein at 4°C overnight. Plates were blocked with 2% nonfat dry milk in DPBS containing 0.05% Tween-20 for 1 h. Each antibody was diluted to a concentration of 10 µg/mL. Next, recombinant human ACE2 protein with a C-terminal FLAG tag was diluted to 2 µg/mL and added to the antibody solution without washing the plate to a final concentration of ACE2 of 0.4µg/mL. ACE2 binding was detected using HRP-conjugated anti-FLAG antibodies and developed with TMB substrate. The reaction was quenched with 1 N hydrochloric acid once color was developed. Absorbance was measured at 450 nm using a spectrophotometer.

### dsRNA staining neutralization assay

Calu-3 cells were seeded at 5,000 cells per well in SCREENSTAR 384-well black plates (Greiner) and allowed to adhere overnight. The cells then were treated with antibodies in 12 concentrations spanning from 5.65 x 10^-5^ µg/mL to 10 µg/mL and immediately transferred to a BSL-3 facility where they were inoculated with SARS-CoV-2 at an approximate MOI of 1 PFU/cell in 50 µL medium, and incubated for 48 h. At the end of the incubation, plates were submerged in PBS with 4% paraformaldehyde and 4% sucrose solution for 30 minutes to fix. Cells then were permeabilized with 0.2% Triton-X-100/PBS for 10 min and blocked with 5% BSA/PBS for 1 h. Primary J2 anti-dsRNA (Scicons #10010500) antibody solution at a 1:1,000 dilution was placed on the cells overnight at 4°C. Cells were washed with 0.1% Tween-20/PBS (PBST) three times and plates were incubated with secondary goat anti-mouse Alexa-Fluor-546-labeled antibody at 1:1,000 dilution (Thermo Fisher Scientific) for 2 h at room temperature in the dark. Plates were washed three times with PBST and incubated with DAPI for 30 min at room temperature in the dark. Plates were then imaged with fluorescent microscopy on a Nikon Eclipse TI2 automated microscopy system with a 20× objective. Six frames per well were imaged and sum dsRNA fluorescence intensity, normalized to cell count by DAPI, was measured by Nikon Elements imaging software.

### Mouse experiments

Animal studies were carried out in accordance with the recommendations in the Guide for the Care and Use of Laboratory Animals of the National Institutes of Health. The protocols were approved by the Institutional Animal Care and Use Committee at the Washington University School of Medicine (assurance number A3381–01). Virus inoculations were performed under anesthesia that was induced and maintained with ketamine hydrochloride and xylazine, and all efforts were made to minimize animal suffering. Heterozygous K18-hACE c57BL/6J mice (strain: 2B6.Cg-Tg(K18-ACE2)2Prlmn/J) were obtained from The Jackson Laboratory. Animals were housed in groups and fed standard chow diets. One day prior to infection, mice were given a single 200 μg dose of COV2-2351 or COV2-2164 by intraperitoneal injection. Eight- to nine-week-old mice were administered 10^3^ PFU of SARS-CoV-2 by intranasal administration.

### Measurement of viral burden in mouse tissues

Tissues were weighed and homogenized with zirconia beads in a MagNA Lyser instrument (Roche Life Science) in 1,000 μL of DMEM medium supplemented with 2% heat-inactivated FBS. Tissue homogenates were clarified by centrifugation at 10,000 rpm for 5 min and stored at −80°C. RNA was extracted using the MagMax mirVana Total RNA isolation kit (Thermo Fisher Scientific) on the Kingfisher Flex extraction robot (Thermo Fisher Scientific). RNA was reverse transcribed and amplified using the TaqMan RNA-to-CT 1-Step Kit (Thermo Fisher). Reverse transcription was carried out at 48°C for 15 min followed by 2 min at 95°C. Amplification was accomplished over 50 cycles as follows: 95°C for 15 s and 60°C for 1 min. The number of copies of SARS-CoV-2 N gene RNA in samples was determined using a previously published assay^68^. Briefly, a TaqMan assay was designed to target a highly conserved region of the N gene (forward primer: ATGCTGCAATCGTGCTACAA; Reverse primer:m GACTGCCGCCTCTGCTC; Probe: /56-FAM/TCAAGGAAC/ZEN/AACATTGCCAA/3IABkFQ/). This region was included in an RNA standard to allow for copy number determination down to 10 copies per reaction. The reaction mixture contained final concentrations of primers or probe of 500 or 100 nM, respectively.

### Electron microscopy sample and grid preparation, imaging and processing of S6P_ecto_–Fab complexes

Fabs were produced by digesting recombinant chromatography-purified IgGs using resin-immobilized cysteine protease enzyme (FabALACTICA, Genovis). The digestion occurred in 100 mM sodium phosphate and 150 mM NaCl pH 7.2 (PBS) for around 16 h at ambient temperature. To remove cleaved Fc from intact IgG, the digestion mix was incubated with CaptureSelect Fc resin (Genovis) for 30 min at ambient temperature in PBS buffer.

For screening and imaging of negatively-stained SARS-CoV-2 S6P_ecto_ protein in complex with human Fabs, the proteins were incubated at a Fab:S molar ratio of 4:1 for about 1 h at ambient temperature or overnight at 4°C. Approximately 3 μL of the sample at concentrations of about 10 to 15 μg/mL was applied to a glow-discharged grid with continuous carbon film on 400 square mesh copper electron microscopy grids (Electron Microscopy Sciences). The grids were stained with 0.75% uranyl formate^69^. Images were recorded on a Gatan US4000 4k × 4k CCD camera using an FEI TF20 (TFS) transmission electron microscope operated at 200 keV and control with Serial EM. All images were taken at 50,000× magnification with a pixel size of 2.18 Å per pixel in low-dose mode at a defocus of 1.5 to 1.8 μm. The total dose for the micrographs was around 30 e− per Å^2^. Image processing was performed using the cryoSPARC software package. Images were imported, CTF-estimated, and particles were picked. The particles were extracted with a box size of 256 pixels and binned to 128 pixels (pixel size of 4.36 Å/pix) and 2D class averages were performed.

### Epitope mapping of antibodies by alanine scanning

Epitope mapping was performed essentially as described previously ^70^ using SARS-CoV-2 (Wuhan-Hu-1 strain) S protein RBD and S2 shotgun mutagenesis mutation libraries, made using a full-length expression construct for S protein. 184 residues of the RBD (between S residues 335 and 526), and 513 S2 residues (between residues 689 -1247) were mutated individually to alanine, and alanine residues to serine. Mutations were confirmed by DNA sequencing, and clones arrayed in a 384-well plate, one mutant per well. Binding of mAbs to each mutant clone in the alanine scanning library was determined, in duplicate, by high-throughput flow cytometry. A plasmid encoding cDNA for each S protein mutant was transfected into HEK-293T cells and allowed to express for 22 h. Cells were fixed in 4% (v/v) paraformaldehyde (Electron Microscopy Sciences), and permeabilized with 0.1% (w/v) saponin (Sigma-Aldrich) in PBS plus calcium and magnesium (PBS++) before incubation with mAbs diluted in PBS++, 10% normal goat serum (Sigma), and 0.1% saponin. MAb screening concentrations were determined using an independent immunofluorescence titration curve against cells expressing wild-type S protein to ensure that signals were within the linear range of detection. Antibodies were detected using 3.75 μg/mL of Alexa-Fluor-488-labeled secondary antibodies (Jackson ImmunoResearch Laboratories) in 10% normal goat serum with 0.1% saponin. Cells were washed three times with PBS++/0.1% saponin followed by two washes in PBS, and mean cellular fluorescence was detected using a high-throughput Intellicyte iQue flow cytometer (Sartorius). Antibody reactivity against each mutant S protein clone was calculated relative to wild-type S protein reactivity by subtracting the signal from mock-transfected controls and normalizing to the signal from wild-type S-transfected controls. Mutations within clones were identified as critical to the mAb epitope if they did not support reactivity of the test MAb but supported reactivity of other SARS-CoV-2 antibodies. This counter-screen strategy facilitates the exclusion of S protein mutants that are locally misfolded or have an expression defect.

### Cell-surface binding to full-length S protein or S2 domain protein

A plasmid encoding the S protein C-terminus S2 region (starting at residue S685) was transfected into HEK-293T cells arrayed in a 384-well plate and allowed to express for 22 h. Cells transfected with vector alone acted as negative controls. MAbs were screened over a range of concentrations, 4 replicates for each mAb concentration, as described for epitope mapping. Fluorescence values were background subtracted.

### ELISA binding assay for serum analysis

To assess serum reactivity, 384-well microtiter plates were coated with purified recombinant SARS-CoV-2 S6P_ecto_ at 4°c overnight. Plates were blocked with blocking buffer (2% non-fat dry milk and 2% normal goat serum in DPBS containing 0.05% Tween-20) for 1 h. Serum was diluted 1:75 in blocking buffer, and then diluted three-fold serially 15 times, and added to wells. Binding was detected with goat anti-human IgG conjugated with horseradish peroxidase and TMB substrate. The reaction was quenched with 1N hydrochloric acid once color was developed. The absorbance was measured at 450 nm using a spectrophotometer (Biotek).

### Plasmablasts isolation and flow cytometric analysis

Blood was collected into tubes containing heparin. To assess plasmablasts frequency in PBMCs for analytical flow cytometric studies, PBMCs were enriched from whole blood (day 10 after first, and day 7 after second vaccination) using direct PBMCs isolation kit (StemCell Technologies). For singe-cell antibody secretion and paired antibody sequencing studies, plasmablasts were enriched from the whole blood (day 7 after second vaccination) by negative selection using custom direct human plasmablasts isolation kit containing paramagnetic beads and antibodies for negative selection (StemCell Technologies). Enriched cells were stained 30 min on ice in a RoboSep buffer (StemCell Technologies) containing following phenotyping antibodies; anti-CD19-FITC (1:20 dilution, eBioscience), anti-CD27-APC (1:20 dilution), and anti-CD38-PE (1:25 dilution, BD Biosciences), and then analyzed by flow cytometry using an SH800 cell sorter (Sony). A DAPI stain was used as a viability dye to exclude dead cells. Plasmablasts were identified as DAPI-CD19^lo^CD27^hi^CD38^hi^ cells. Approximately 40,000 and ∼6,000 plasmablasts were FACS-sorted in a bulk for paired antibody sequencing and single-cell antibody secretion studies, respectively.

### Generation of antibody variable-gene libraries from single plasmablasts

For paired antibody sequencing, cells were resuspended into DPBS containing 0.04% non-acetylated BSA, split into four replicates, and separately added to 50 μL of RT Reagent Mix, 5.9 μL of Poly-dt RT Primer, 2.4 μL of Additive A and 10 μL of RT Enzyme Mix B to complete the Reaction Mix as per the vendor’s protocol. The reactions then were loaded onto a Chromium chip (10x Genomics). Chromium Single Cell V(D)J B-Cell-enriched libraries were generated, quantified, normalized and sequenced according to the User Guide for Chromium Single Cell V(D)J Reagents kits (CG000086_REV C). Amplicons were sequenced on an Illumina Novaseq 6000, and data were processed using the CellRanger software v3.1.0 (10X Genomics).

### Single-cell antibody secretion analysis using Beacon instrument

FACS-purified plasmablasts were resuspended in plasmablast survival medium that promotes antibody secretion and assessed for reactivity of secreted antibodies using the 11k chip on Beacon optofluidic instrument (Berkley Lights) as previously described^8^. Single cell-antibody secretion binding assay was performed as previously described^8^ using SARS-CoV-2 S6P_ecto_- and SARS-CoV-2 RBD-coated beads.

### ELISpot assay

Direct enzyme-linked immunosorbent spot (ELISpot) assay was performed to enumerate plasmablasts present in the PBMC samples secreting IgG, IgM, or IgA antibodies reacting with either SARS-CoV-2-S6P_ecto_ protein or influenza A/Darwin/42/2020 H1N1 hemagglutinin protein (as a negative control). Briefly, 96-well ELISpot MSIP plates (Millipore) were activated with 100 µL 100% methanol/well for 10 sec, washed three times with 1× DPBS, coated overnight either with 100 µL of 2 µg/mL of SARS-CoV-2-S6P_ecto_ or influenza HA protein in PBS overnight at 4°C. Plates were washed three times with 1× DPBS and blocked by incubation with RPMI containing 10% FCS at 37°C for 2 h. Enriched plasmablasts or FACS-sorted plasmablasts were added to the plates and incubated 18-24 h at 37°C. Plates were washed with PBS and then PBS containing 0.05% Tween, and then incubated with either goat anti-human IgG-HRP conjugated antibodies (Southern Biotech), goat anti-human IgA-HRP conjugated antibodies (Southern Biotech), or goat anti-human IgM-HRP conjugated antibodies (Southern Biotech) for 2 h at room temperature. After washing three times with PBS containing 0.05% Tween/1% BSA, plates were developed using 3-amino-9-ethyl-carbazole (AEC) substrate (Sigma). The developed plates were scanned, and spots were analyzed using an automated ELISpot counter (Cellular Technologies Ltd.). Plasmablasts or sorted plasmablasts from PBMCs were added to the plates and incubated 18-24 h at 37°C. Plates were washed with PBS and then PBS containing 0.05% Tween, and then incubated with either goat anti-human IgG-HRP conjugated antibodies (Southern Biotech, catalog no. 2040-05), goat anti-human IgA-HRP conjugated antibodies (Southern Biotech, catalog no. 2050-05), or goat anti-human IgM-HRP conjugated antibodies (Southern Biotech, catalog no. 2020-05) for 2 h at room temperature. After washing three times with PBS containing 0.05% Tween/1% BSA, plates were developed using 3-amino-9-ethyl-carbazole (AEC) substrate (Sigma). The developed plates were scanned and spots were analyzed using an automated ELISpot counter (Cellular Technologies Ltd.).

### Quantification and statistical analysis

The descriptive statistics mean ± SEM or mean ± SD were determined for continuous variables as noted. Virus titers in the tissues were compared using one-way ANOVA with Turkey’s post-test. Curves for antibody binding and neutralization were fitted after log transformation of antibody concentrations using non-linear regression analysis. Technical and biological replicates are indicated in the figure legends. Statistical analyses were performed using Prism v8.4.3 (GraphPad).

## References

1. Jiang, S., Hillyer, C. & Du, L. Neutralizing antibodies against SARS-CoV-2 and other human coronaviruses. Trends Immunol 41, 355–359, doi:10.1016/j.it.2020.03.007 (2020).

2. Krammer, F. SARS-CoV-2 vaccines in development. Nature 586, 516–527, doi:10.1038/s41586-020-2798-3 (2020).

3. Bosch, B. J., van der Zee, R., de Haan, C. A. & Rottier, P. J. The coronavirus spike protein is a class I virus fusion protein: structural and functional characterization of the fusion core complex. J Virol 77, 8801–8811, doi:10.1128/jvi.77.16.8801-8811.2003 (2003).

4. Tortorici, M. A. & Veesler, D. Structural insights into coronavirus entry. Adv Virus Res 105, 93–116, doi:10.1016/bs.aivir.2019.08.002 (2019).

5. Wan, Y., Shang, J., Graham, R., Baric, R. S. & Li, F. Receptor recognition by the novel coronavirus from Wuhan: an analysis based on decade-long structural studies of SARS coronavirus. J Virol 94, doi:10.1128/JVI.00127-20 (2020).

6. Hoffmann, M. et al. SARS-CoV-2 cell entry depends on ACE2 and TMPRSS2 and is blocked by a clinically proven protease inhibitor. Cell 181, 271–280 e278, doi:10.1016/j.cell.2020.02.052 (2020).

7. Li, W. et al. Angiotensin-converting enzyme 2 is a functional receptor for the SARS coronavirus. Nature 426, 450–454, doi:10.1038/nature02145 (2003).

8. Zost, S. J. et al. Rapid isolation and profiling of a diverse panel of human monoclonal antibodies targeting the SARS-CoV-2 spike protein. Nature Medicine, doi:10.1038/s41591-020-0998-x (2020).

9. Robbiani, D. F. et al. Convergent antibody responses to SARS-CoV-2 infection in convalescent individuals. bioRxiv, doi:10.1101/2020.05.13.092619 (2020).

10. Liu, L. et al. Potent neutralizing antibodies against multiple epitopes on SARS-CoV-2 spike. Nature 584, 450–456, doi:10.1038/s41586-020-2571-7 (2020).

11. Seydoux, E. et al. Characterization of neutralizing antibodies from a SARS-CoV-2 infected individual. bioRxiv, doi:10.1101/2020.05.12.091298 (2020).

12. Wec, A. Z. et al. Broad neutralization of SARS-related viruses by human monoclonal antibodies. Science 369, 731–736, doi:10.1126/science.abc7424 (2020).

13. Regeneron Pharmaceuticals, Inc. “Regeneron’s casirivimab and imdevimab antibody cocktail for COVID-19 is first combination to receive FDA emergency use authorization,” press release (21 November, 2020); https://investor.regeneron.com/news-releases/news-release-details/regenerons-regen-cov2-first-antibody-cocktail-covid-19-receive/.

14. Eli Lilly and Company, “ A phase 3 randomized, double-blind, placebo-controlled trial to evaluate the efficacy and safety of LY3819253 alone and in combination with LY3832479 in preventing SARS-CoV-2 infection and COVID-19 in skilled nursing and assisted living facility residents and staff; a NIAID and Lilly Collaborative Study” (Clinical trial registration NCT04497987, clinicaltrials.gov, 2020); https://clinicaltrials.gov/ct2/show/NCT04497987.

15. Johnson and Johnson, “Johnson & Johnson Announces U.S. CDC Advisory Committee Recommends First Single-Shot COVID-19 Vaccine for Adults 18 and Older in U.S.”, press release (February 28th, 2021); https://www.jnj.com/johnson-johnson-announces-u-s-cdc-advisory-committee-recommends-first-single-shot-covid-19-vaccine-for-adults-18-and-older-in-u-s.

16. Pfizer, “Pfizer and BioNTech announch vaccine candidate against COVID-19 achieved success in first interim analysis from phase 3 study”, press release (November 09, 2020); https://www.pfizer.com/news/press-release/press-release-detail/pfizer-and-biontech-announce-vaccine-candidate-against.

17. Moderna,“Moderna’s COVID-19 vaccine candidate meets its primary efficacy endpoint in the first interim analysis of the phase 2 COVE study”, press release (November 16, 2020); https://investors.modernatx.com/news-releases/news-release-details/modernas-covid-19-vaccine-candidate-meets-its-primary-efficacy.

18. Cohen-Dvashi, H. et al. Structural basis for a convergent immune response against Ebola virus. Cell Host Microbe 27, 418–427 e414, doi:10.1016/j.chom.2020.01.007 (2020).

19. Ehrhardt, S. A. et al. Polyclonal and convergent antibody response to Ebola virus vaccine rVSV-ZEBOV. Nat Med 25, 1589–1600, doi:10.1038/s41591-019-0602-4 (2019).

20. Davis, C. W. et al. Longitudinal analysis of the human B cell response to Ebola virus infection. Cell 177, 1566–1582 e1517, doi:10.1016/j.cell.2019.04.036 (2019).

21. Seth J. Zost, J. D., Iuliia Gilchuk, Pavlo Gilchuk, Natalie J. Thornburg, Sandhya Bangaru, Nurgun Kose, Jessica A. Finn, Robin Bombardi, Cinque Soto, Rachel Nargi, Ryan Irving, Naveenchandra Suryadevara, Jonna B. Westover, Robert H. Carnahan, Hannah L. Turner, Sheng Li, Andrew B. Ward, James E. Crowe Jr. . Canonical features of human antibodies recognizing the influenza hemagglutinin trimer interface. bioRxiv, doi:https://doi.org/10.1101/2020.12.31.4_24868 (2021).

22. Pappas, L. et al. Rapid development of broadly influenza neutralizing antibodies through redundant mutations. Nature 516, 418–422, doi:10.1038/nature13764 (2014).

23. Joyce, M. G. et al. Vaccine-induced antibodies that neutralize Group 1 and Group 2 influenza A viruses. Cell 166, 609–623, doi:10.1016/j.cell.2016.06.043 (2016).

24. Sui, J. et al. Structural and functional bases for broad-spectrum neutralization of avian and human influenza A viruses. Nat Struct Mol Biol 16, 265–273, doi:10.1038/nsmb.1566 (2009).

25. Wheatley, A. K. et al. H5N1 vaccine-elicited memory B cells are genetically constrained by the IGHV locus in the recognition of a neutralizing epitope in the hemagglutinin stem. J Immunol 195, 602–610, doi:10.4049/jimmunol.1402835 (2015).

26. Setliff, I. et al. Multi-donor longitudinal antibody repertoire sequencing reveals the existence of public antibody clonotypes in HIV-1 infection. Cell Host Microbe 23, 845–854 e846, doi:10.1016/j.chom.2018.05.001 (2018).

27. Wu, X. et al. Focused evolution of HIV-1 neutralizing antibodies revealed by structures and deep sequencing. Science 333, 1593–1602, doi:10.1126/science.1207532 (2011).

28. Zhou, T. et al. Structural repertoire of HIV-1-neutralizing antibodies targeting the CD4 supersite in 14 donors. Cell 161, 1280–1292, doi:10.1016/j.cell.2015.05.007 (2015).

29. Williams, W. B. et al. HIV-1 VACCINES. Diversion of HIV-1 vaccine-induced immunity by gp41-microbiota cross-reactive antibodies. Science 349, aab1253, doi:10.1126/science.aab1253 (2015).

30. Bailey, J. R. et al. Broadly neutralizing antibodies with few somatic mutations and hepatitis C virus clearance. JCI Insight 2, doi:10.1172/jci.insight.92872 (2017).

31. Giang, E. et al. Human broadly neutralizing antibodies to the envelope glycoprotein complex of hepatitis C virus. Proc Natl Acad Sci U S A 109, 6205–6210, doi:10.1073/pnas.1114927109 (2012).

32. Yuan, M. et al. Structural basis of a shared antibody response to SARS-CoV-2. Science 369, 1119–1123, doi:10.1126/science.abd2321 (2020).

33. Dong, J. et al. Genetic and structural basis for recognition of SARS-CoV-2 spike protein by a two-antibody cocktail. bioRxiv, doi:10.1101/2021.01.27.428529 (2021).

34. Nielsen, S. C. A. et al. B cell clonal expansion and convergent antibody responses to SARS-CoV-2. Res Sq, doi:10.21203/rs.3.rs-27220/v1 (2020).

35. Soto, C. et al. High frequency of shared clonotypes in human B cell receptor repertoires. Nature 566, 398–402, doi:10.1038/s41586-019-0934-8 (2019).

36. Briney, B., Inderbitzin, A., Joyce, C. & Burton, D. R. Commonality despite exceptional diversity in the baseline human antibody repertoire. Nature 566, 393–397, doi:10.1038/s41586-019-0879-y (2019).

37. Tan, T. J. C. et al. Sequence signatures of two IGHV3-53/3-66 public clonotypes to SARS-CoV-2 receptor binding domain. bioRxiv, doi:10.1101/2021.01.26.428356 (2021).

38. Brouwer, P. J. M. et al. Potent neutralizing antibodies from COVID-19 patients define multiple targets of vulnerability. Science 369, 643–650, doi:10.1126/science.abc5902 (2020).

39. Kreer, C. et al. Longitudinal isolation of potent near-germline SARS-CoV-2-neutralizing antibodies from COVID-19 patients. Cell 182, 843–854 e812, doi:10.1016/j.cell.2020.06.044 (2020).

40. Rogers, T. F. et al. Isolation of potent SARS-CoV-2 neutralizing antibodies and protection from disease in a small animal model. Science 369, 956–963, doi:10.1126/science.abc7520 (2020).

41. Cao, Y. et al. Potent Neutralizing Antibodies against SARS-CoV-2 Identified by High-Throughput Single-Cell Sequencing of Convalescent Patients’ B Cells. Cell 182, 73–84 e16, doi:10.1016/j.cell.2020.05.025 (2020).

42. Zost, S. J. et al. Potently neutralizing and protective human antibodies against SARS-CoV-2. Nature 584, 443–449, doi:10.1038/s41586-020-2548-6 (2020).

43. Gilchuk, P. et al. Integrated pipeline for the accelerated discovery of antiviral antibody therapeutics. Nat Biomed Eng 4, 1030–1043, doi:10.1038/s41551-020-0594-x (2020).

44. Ladner, J. T. et al. Epitope-resolved profiling of the SARS-CoV-2 antibody response identifies cross-reactivity with endemic human coronaviruses. Cell Rep Med 2, 100189, doi:10.1016/j.xcrm.2020.100189 (2021).

45. Hansen, J. et al. Studies in humanized mice and convalescent humans yield a SARS-CoV-2 antibody cocktail. Science 369, 1010–1014, doi:10.1126/science.abd0827 (2020).

46. Fibriansah, G. et al. DENGUE VIRUS. Cryo-EM structure of an antibody that neutralizes dengue virus type 2 by locking E protein dimers. Science 349, 88–91, doi:10.1126/science.aaa8651 (2015).

47. Yuan, M. et al. A highly conserved cryptic epitope in the receptor binding domains of SARS-CoV-2 and SARS-CoV. Science 368, 630–633, doi:10.1126/science.abb7269 (2020).

48. Winkler, E. S. et al. SARS-CoV-2 infection of human ACE2-transgenic mice causes severe lung inflammation and impaired function. Nat Immunol 21, 1327–1335, doi:10.1038/s41590-020-0778-2 (2020).

49. Zheng, J. et al. COVID-19 treatments and pathogenesis including anosmia in K18-hACE2 mice. Nature 589, 603–607, doi:10.1038/s41586-020-2943-z (2021).

50. Golden, J. W. et al. Human angiotensin-converting enzyme 2 transgenic mice infected with SARS-CoV-2 develop severe and fatal respiratory disease. JCI Insight 5, doi:10.1172/jci.insight.142032 (2020).

51. Micah Rapp, Y. G., Eswar R. Reddem, Lihong Liu, Pengfei Wang, Jian Yu, Gabriele Cerutti, Jude Bimela, Fabiana Bahna, Seetha Mannepalli, Baoshan Zhang, Peter D. Kwong, David D. Ho, Lawrence Shapiro, Zizhang Sheng. Modular basis for potent SARS-CoV-2 neutralization by a prevalent VH1-2-derived antibody class. bioRxiv (2021).

52. Tortorici, M. A. et al. Ultrapotent human antibodies protect against SARS-CoV-2 challenge via multiple mechanisms. Science 370, 950–957, doi:10.1126/science.abe3354 (2020).

53. Voss, W. N. et al. Prevalent, protective, and convergent IgG recognition of SARS-CoV-2 non-RBD spike epitopes in COVID-19 convalescent plasma. bioRxiv, doi:10.1101/2020.12.20.423708 (2020).

54. Yuan, M. et al. Structural and functional ramifications of antigenic drift in recent SARS-CoV-2 variants. bioRxiv, doi:10.1101/2021.02.16.430500 (2021).

55. Wibmer, C. K. et al. SARS-CoV-2 501Y.V2 escapes neutralization by South African COVID-19 donor plasma. bioRxiv, doi:10.1101/2021.01.18.427166 (2021).

56. Linda J. Rennick, L. R. R.-M., Sham Nambulli, W. Paul Duprex, Kevin R. McCarthy. Deletion disrupts a conserved antibody epitope in a SARS-CoV-2 variant of concern. bioRxiv (2021).

57. Cerutti, G. et al. Potent SARS-CoV-2 Neutralizing Antibodies Directed Against Spike N-Terminal Domain Target a Single Supersite. Cell Host & Microbe, doi:10.1016/j.chom.2021.03.005.

58. Suryadevara, N. et al. Neutralizing and protective human monoclonal antibodies recognizing the N-terminal domain of the SARS-CoV-2 spike protein. bioRxiv, 2021.2001.2019.427324, doi:10.1101/2021.01.19.427324 (2021).

59. Rappuoli, R., Bottomley, M. J., D’Oro, U., Finco, O. & De Gregorio, E. Reverse vaccinology 2.0: Human immunology instructs vaccine antigen design. J Exp Med 213, 469–481, doi:10.1084/jem.20151960 (2016).

60. Anderson, E. M. et al. Seasonal human coronavirus antibodies are boosted upon SARS-CoV-2 infection but not associated with protection. Cell, doi:10.1016/j.cell.2021.02.010 (2021).

61. Collier, D. A. et al. Sensitivity of SARS-CoV-2 B.1.1.7 to mRNA vaccine-elicited antibodies. Nature, doi:10.1038/s41586-021-03412-7 (2021).

62. Wang, P. et al. Antibody resistance of SARS-CoV-2 variants B.1.351 and B.1.1.7. Nature, doi:10.1038/s41586-021-03398-2 (2021).

63. Tegally, H. et al. Emergence of a SARS-CoV-2 variant of concern with mutations in spike glycoprotein. Nature, doi:10.1038/s41586-021-03402-9 (2021).

64. Wang, Z. et al. mRNA vaccine-elicited antibodies to SARS-CoV-2 and circulating variants. Nature, doi:10.1038/s41586-021-03324-6 (2021).

65. Case, J. B. et al. Neutralizing antibody and soluble ACE2 inhibition of a replication-competent VSV-SARS-CoV-2 and a clinical isolate of SARS-CoV-2. Cell Host Microbe 28, 475–485 e475, doi:10.1016/j.chom.2020.06.021 (2020).

66. Soto, C. et al. PyIR: a scalable wrapper for processing billions of immunoglobulin and T cell receptor sequences using IgBLAST. BMC Bioinformatics 21, 314, doi:10.1186/s12859-020-03649-5 (2020).

67. Chng, J. et al. Cleavage efficient 2A peptides for high level monoclonal antibody expression in CHO cells. MAbs 7, 403–412, doi:10.1080/19420862.2015.1008351 (2015).

68. Case, J. B., Bailey, A. L., Kim, A. S., Chen, R. E. & Diamond, M. S. Growth, detection, quantification, and inactivation of SARS-CoV-2. Virology 548, 39–48, doi:10.1016/j.virol.2020.05.015 (2020).

69. Ohi, M., Li, Y., Cheng, Y. & Walz, T. Negative staining and image classification - Powerful tools in modern electron microscopy. Biol Proced Online 6, 23–34, doi:10.1251/bpo70 (2004).

70. Davidson, E. & Doranz, B. J. A high-throughput shotgun mutagenesis approach to mapping B-cell antibody epitopes. Immunology 143, 13–20, doi:10.1111/imm.12323 (2014).

