## Supplemental Figures for "Convergent antibody responses to the SARS-CoV-2 spike protein in convalescent and vaccinated individuals"

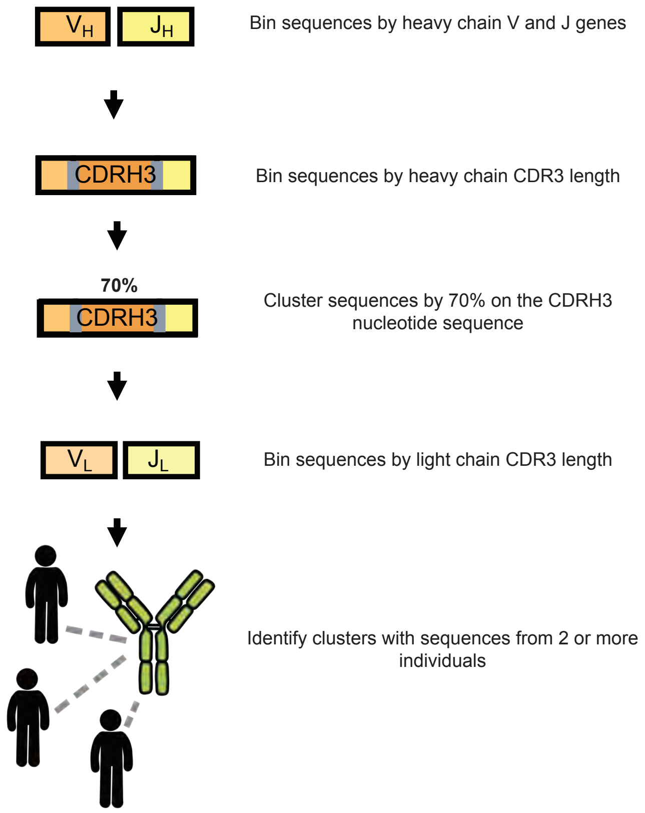
VH
JH
Bin sequences by heavy chain V and J genes
CDRH3
Bin sequences by heavy chain CDR3 length
70%
CDRH3
Cluster sequences by 70% on the CDRH3 nucleotide sequence
VL
JL
Bin sequences by light chain CDR3 length
Identify clusters with sequences from 2 or more individuals

### Extended data Fig. 2

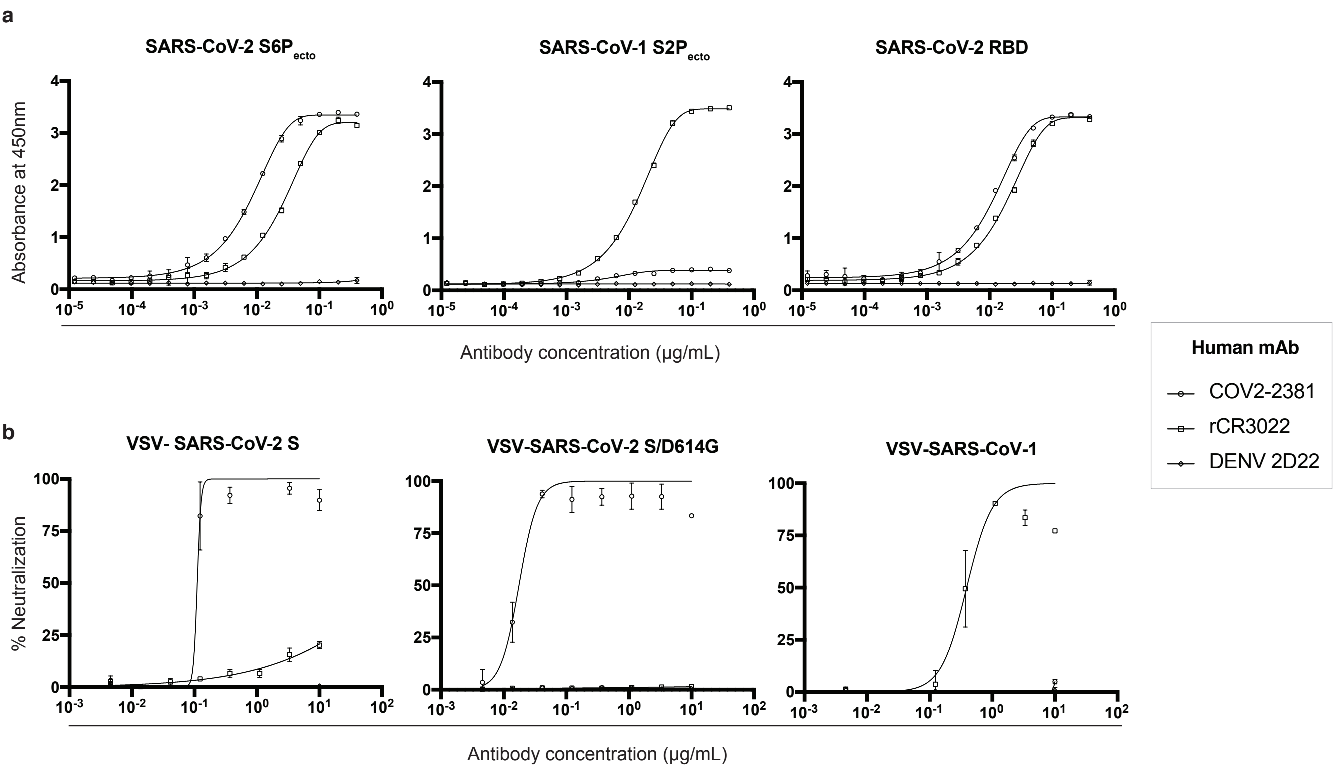

#### Extended data Fig. 3

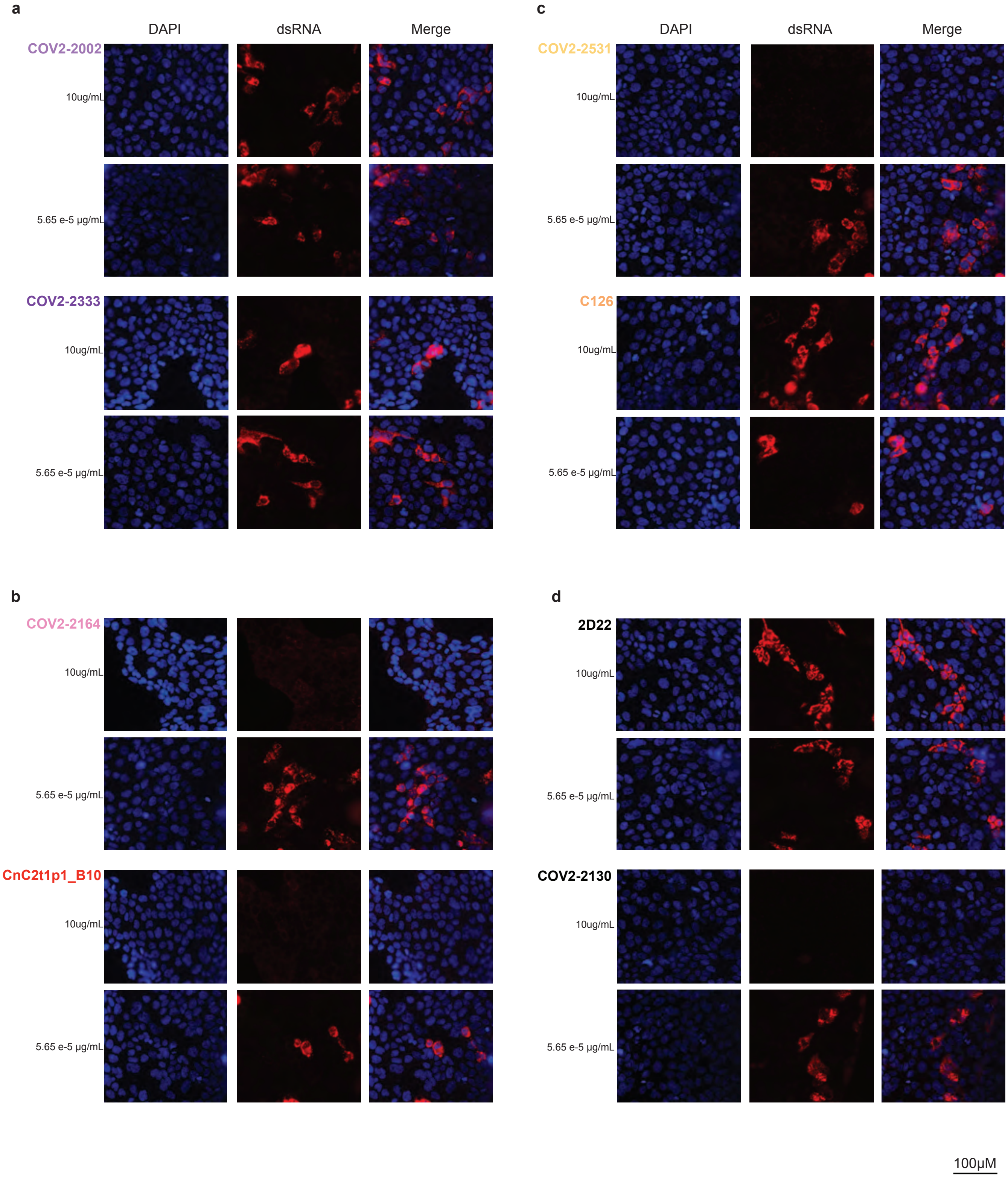

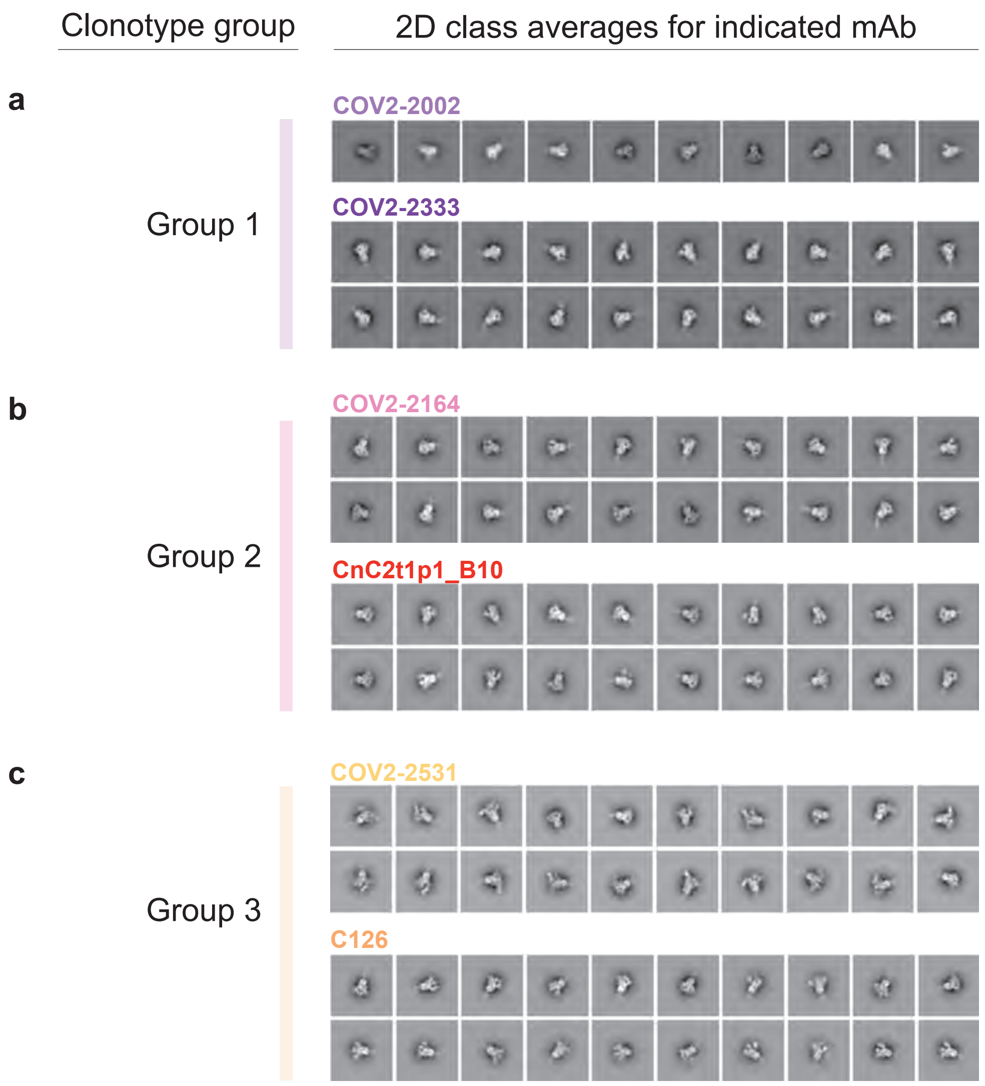
Clonotype group
2D class averages for indicated mAb
a
Group 1
COV2-2002
COV2-2333
b
Group 2
COV2-2164
CnC2t1p1_B10
c
Group 3
COV2-2531
C126

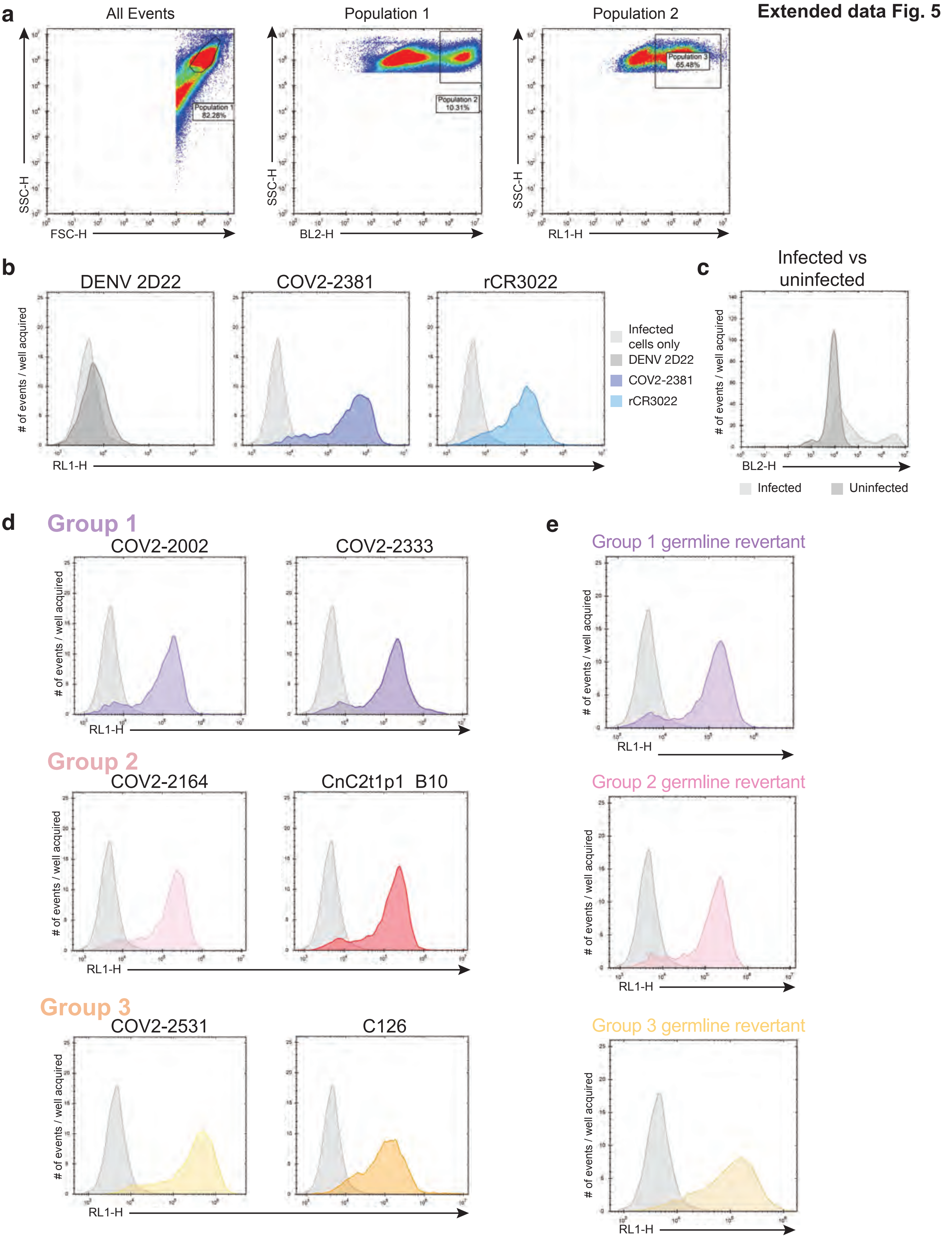

Extended data Fig. 5
a
All Events
Population 1
Population 2
Population 1 82.28%
Population 2 10.31%
Population 3 65.48%
SSC-H
FSC-H
BL2-H
RL1-H
b
DENV 2D22
COV2-2381
rCR3022
Infected cells only
DENV 2D22
COV2-2381
rCR3022
### of events / well acquired
RL1-H
c
Infected vs uninfected
### of events / well acquired
BL2-H
Infected
Uninfected
d
Group 1
COV2-2002
COV2-2333
Group 2
COV2-2164
CnC2t1p1 B10
Group 3
COV2-2531
C126
### of events / well acquired
RL1-H
e
Group 1 germline revertant
Group 2 germline revertant
Group 3 germline revertant
### of events / well acquired
RL1-H

**a**

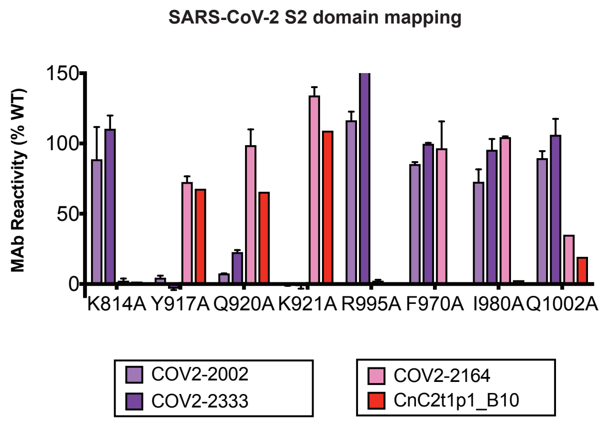
SARS-CoV-2 S2 domain mapping
MAb Reactivity (% WT)
150
100
50
0
K814A Y917A Q920A K921A R995A F970A I980A Q1002A
COV2-2002
COV2-2333
COV2-2164
CnC2t1p1_B10

**b**

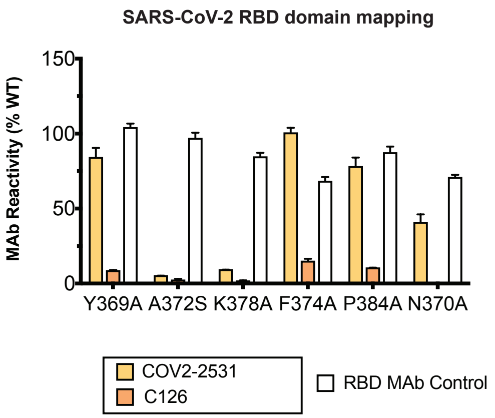
SARS-CoV-2 RBD domain mapping
MAb Reactivity (% WT)
150
100
50
0
Y369A A372S K378A F374A P384A N370A
COV2-2531
C126
RBD MAb Control

**Extended data Fig. 7**

| Virus | Amino acid sequence of the RBD protein, at the indicated positions | | |
|---|---|---|---|
| SARS-CoV-1 | 306 | RVVPSGDVVRFPNITNLCPFGEVFNATKFPSVYAWERKKISNCVADYSVL | 355 |
| SARS-CoV-2 | 319 | RVQPTESIVRFPNITNLCPFGEVFNATRFASVYAWNRKRISNCVADYSVL | 368 |
| SARS-CoV-1 | 356 | YNSTFFSTFKCYGVSATKLNDLCFSNVYADSFVVKGDDVRQIAPGQTGVI | 405 |
| SARS-CoV-2 | 369 | YNSASFSTFKCYGVSPTKLNDLCFTNVYADSFVIRGDEVRQIAPGQTGKI | 418 |
| SARS-CoV-1 | 406 | ADYNYKLPDDFMGCVLAWNTRNIDATSTGNHNYKYRYLRHGKLRPFERDI | 455 |
| SARS-CoV-2 | 419 | ADYNYKLPDDFTGCVIAWNSNNLDSKVGGNYNYLYRLFRKSNLKPFERDI | 468 |
| SARS-CoV-1 | 456 | SNVPFSPDGKPCTP-PALNCYWPLNDYGFYTTTGIGYQPYRVVVLSFELL | 504 |
| SARS-CoV-2 | 469 | STEIYQAGSTPCNGVEGFNCYFPLQSYGFQPTNGVGYQPYRVVVLSFELL | 518 |
| SARS-CoV-1 | 505 | NAPATVCGPKLSTDLIKNQCVNF | |
| SARS-CoV-2 | 519 | HAPATVCGPKKSTNLVKNKCVNF | |

CR3022 epitope

COV2-2531 critical residues

C126 critical residues

#### Extended data Fig. 8

**Neutralizing potency**

| Variability of epitope | High | Low |
|---|---|---|
| Low | **Class 1**<br>Neutralizing<br>Invariant residues<br>ex.: *IGHV1-58* | **Class 3**<br>Non-neutralizing<br>Invariant residues<br>ex.: *IGHV3-7* |
| High | **Class 2**<br>Neutralizing<br>Variable residues<br>ex.: *IGHV3-53, 3-66* | **Class 4**<br>Non-neutralizing<br>Variable residues |
